# The developmental hourglass model is applicable to the spinal cord

**DOI:** 10.1101/2021.03.08.434342

**Authors:** Katsuki Mukaigasa, Chie Sakuma, Hiroyuki Yaginuma

**Affiliations:** Department of Neuroanatomy and Embryology, School of Medicine, Fukushima Medical University

**Keywords:** The developmental hourglass model, Single cell RNA-seq, *Cis* regulatory module, Gene regulatory network, Evolution, Spinal cord

## Abstract

The developmental hourglass model predicts that embryonic morphology is most conserved at the mid-embryonic stage and diverges at the early and late stages. This model is generally considered by whole embryonic level. Here, we demonstrate that the hourglass model is also applicable to a reduced element, the spinal cord. In the middle of spinal cord development, dorsoventrally arrayed neuronal progenitor domains are established, which are conserved among vertebrates. We found that, by comparing the single-cell transcriptomes between mice and zebrafish, V3 interneurons, a subpopulation of the post-mitotic spinal neurons, display divergent molecular profiles. We also found non-conservation of *cis*-regulatory elements located around the progenitor fate determinants, indicating the rewiring of the upstream gene regulatory network. These results demonstrate that, despite the conservation of the progenitor domains, processes before and after the progenitor domain specification diverged. This study may help understand the molecular basis of the developmental hourglass model.

## Introduction

The vertebrate neural tube can produce a diverse array of neurons in a precisely controlled and reproducible manner. During neurogenesis and neuronal differentiation, each neuron is assigned a distinct feature, such as neurotransmitter phenotype, axonal projection pathway, and cell body localization (Lai et al., 2016; Lu et al., 2015; Sagner and Briscoe, 2019). The initial phase of this process is the fate specification of progenitor cells, whereby molecularly defined progenitor domains are established with sharp boundaries along the dorsoventral axis of the neural tube (Briscoe et al., 2000). Each progenitor identity is specified through a highly complex gene regulatory network (GRN), which is composed of graded Shh signaling activity localized ventrally, Bmp and Wnt signaling molecules expressed dorsally, pan-neural transcriptional activator Sox1–3, and a number of domain-specific transcription factors (TFs) functioning as repressors (Andrews et al., 2019; Balaskas et al., 2012; Delás and Briscoe, 2020; Kutejova et al., 2016; Nishi et al., 2015; Oosterveen et al., 2012; Peterson et al., 2012; Sagner and Briscoe, 2019; Zagorski et al., 2017). As the output of the GRN, 11 progenitor domains, termed dp1–6 in the dorsal half, and p0, p1, p2, pMN and p3 in the ventral half, are established (Lai et al., 2016; Lu et al., 2015; Sagner and Briscoe, 2019). Subsequently, multiple neuronal subtypes are generated from a single progenitor domain. For example, V2a, V2b and V2c interneurons (INs) are differentiated from the p2 domain (Del Barrio et al., 2007; Li et al., 2005; Panayi et al., 2010; Peng et al., 2007), and dI1i and dI1c INs are differentiated from the dp1 domain (Ding et al., 2012; Wilson et al., 2008). Each subtype of post-mitotic neurons proceeds to the maturation process, such as migration, axonal projection and circuit formation.

These knowledges have been constructed mainly based on the chick and mouse studies, while a number of studies using zebrafish have demonstrated that the spatial architecture of the progenitor domains in the neural tube is largely conserved among vertebrates (Cheesman and Eisen, 2004; Cheesman et al., 2004; Gribble et al., 2007; Guner and Karlstrom, 2007; Lewis et al., 2005; Park et al., 2002; Schäfer et al., 2005).

However, there exist overt differences between amniotes and teleosts with regard to post-mitotic neuronal maturation. For example, V2a INs, which are defined by the expression of Vsx2 (Chx10), contribute to the locomotor rhythm generation in zebrafish (Eklöf-Ljunggren et al., 2012), whereas V2a INs in mice play a role in the left-right alternation of limbs by providing excitatory input to the commissural INs (Crone et al., 2008, 2009). This implies that, during post-mitotic differentiation in mice and zebrafish, V2a INs are assigned different properties, or are placed in different positions within the spinal locomotor circuit (Kiehn, 2016). Another example is Robo3, an axon guidance receptor that is essential for commissural axons to cross the midline (Friocourt and Chédotal, 2017; Marillat et al., 2004; Sabatier et al., 2004). In the spinal cord of amniotes, Robo3 is expressed in V0, V1, and V3 INs in the ventral spinal cord, which encompass commissural INs (Friocourt et al., 2019; Tulloch et al., 2019). However, in zebrafish, although double labeling of *robo3* and neuronal subtype markers is not conducted, *robo3* expression is observed in the region encompassing MNs (Challa et al., 2005). If zebrafish MNs express *bona fide robo3*, the role of Robo3 has diverged between amniotes and zebrafish, as MNs of amniotes never express Robo3 (Friocourt et al., 2019; Tulloch et al., 2019).

In addition to the post-mitotic differentiation, a process before the establishment of progenitor domains also differs in amniotes and zebrafish. *Shh* is an essential gene for specification of the ventral neural tube, and is expressed in the notochord and floor plate in amniotes (Echelard et al., 1993; Riddle et al., 1993; Roelink et al., 1994). In contrast, 3 *Shh*-related genes, *shha* (*sonic hedgehog*), *shhb* (*tiggy winkle hedgehog*) and *ihhb* (*echidna hedgehog*) are expressed in notochord and/or floor plate in zebrafish, conferring a functional redundancy of hedgehog (Hh) signaling (Currie and Ingham, 1996; Ekker et al., 1995; Krauss et al., 1993). The Hh signaling pathway culminates in Gli family TFs, which function as transcriptional activator or repressor depending on Hh signaling activity (Briscoe and Thérond, 2013). Amniotes possess three *Gli* genes (*Gli1*, *Gli2* and *Gli3*), whereas teleosts possess four *Gli* genes (*gli1*, *gli2a*, *gli2b* and *gli3*). Loss of function experiments of *Gli* genes led to clearly different phenotypes between mice and zebrafish. For example, *Gli1* knock-out mice are viable and appear normal, showing that *Gli1* is not essential for embryogenesis in mice (Park et al., 2000). In contrast, *gli1* mutant zebrafish displayed severe cranial motor neuron deficiency and reduced Hh-target genes such as *ptch1* and *nkx2.2a*, and died at the larval stage (Chandrasekhar et al., 1999; Karlstrom et al., 1996, 2003). In *Gli2* knock-out mice, the floor plate was not specified, and MNs aberrantly occupied the ventral-most domain in the spinal cord, though MN itself was differentiated (Ding et al., 1998; Matise et al., 1998), whereas in zebrafish, *gli2a* is completely dispensable for normal embryogenesis and growth to adulthood (Karlstrom et al., 2003; Vanderlaan et al., 2005; Wang et al., 2013), and *gli2b* knock-down caused a marked reduction of MNs in the spinal cord (Ke et al., 2008). Knock-down of *gli3* in zebrafish led to the reduction of MNs (Vanderlaan et al., 2005); however, in *Gli3* mutant mice, patterning defects of the floor plate and MNs were not observed (Persson et al., 2002).

Overall, the processes before and after the progenitor domain specification have diverged among vertebrates, despite the conservation of the dorsoventral arrangement of the progenitor domains. This pattern of developmental divergence is highly consistent with the developmental hourglass model, which argue that embryonic morphology at the early and late developmental stages are divergent and that that at the mid-embryonic stage is conserved (Duboule, 1994; Hu et al., 2017; Irie and Kuratani, 2014). However, the process of spinal cord development has never been investigated from the perspective of this model. Here, we provide evidence supporting that the developmental hourglass model is applicable to spinal cord development. We examined the single-cell transcriptome data from mice and zebrafish, and provide another example of diverse differentiation of post-mitotic neurons. We also examined the transcriptional regulatory elements in the neural tube patterning genes based on ChIP-seq and ATAC-seq data, and sequence conservation, suggesting that GRN upstream to the progenitor domain specification has been rewired during vertebrate evolution. Based on our findings and robust development of the neuronal progenitor specification (Balaskas et al., 2012; Delás and Briscoe, 2020; Exelby et al., 2021; Xiong et al., 2013; Zagorski et al., 2017), we propose that the progenitor domain configuration in the neural tube is less evolvable due to its canalization (Waddington, 1942, 1957).

## Results

### Two distinct subtypes of V3 INs in amniotes

Neuronal subtypes repertoire of V3 INs, which are differentiated from ventral-most p3 progenitor domain, is relatively less studied among spinal neurons, both in amniotes and teleosts. Thus, we focused on V3 INs to clarify to what extent the post mitotic neuronal differentiation has diverged among vertebrates. We examined single-cell RNA-sequence (scRNA-seq) data obtained from the spinal cord of mouse embryos (Delile et al., 2019). The data consist of the whole spinal cord cells from embryonic day (E) 9.5 to E13.5. We extracted data with V3 IN profiles (*Nkx2-2* or *Sim1* positive), applied graph-based clustering and dimensionality reduction by tSNE to these cells, and then identified the differentially expressed genes in each cluster (Table S1). The expression levels of several markers were visualized on tSNE plots (Fig.1A). *Nkx2-2* is a marker of p3 progenitor domain and V3 INs (Briscoe et al., 1999). *Sim1* is a post-mitotic V3 IN marker (Zhang et al., 2008). V3 INs are glutamatergic neurons, and thus express *Slc17a6* (*vGlut2*) (Zhang et al., 2008). Expression of *Sox2*, *Neurog3* and *Tubb3* (*class III β-tubulin*) in this order indicates the transition from progenitor to differentiated neuron (Carcagno et al., 2014). Thus, medial to lateral direction in the spinal cord corresponds to top to bottom direction in this plot. We found at least two distinct populations in V3 INs; one positive for *Robo3*, *Olig3*, and *Cntn2* (*Tag1*) (Cluster 5 in Fig.1A), and the other positive for *Lhx1* (Cluster 1 in Fig.1A, Table S1).

**Figure 1.**
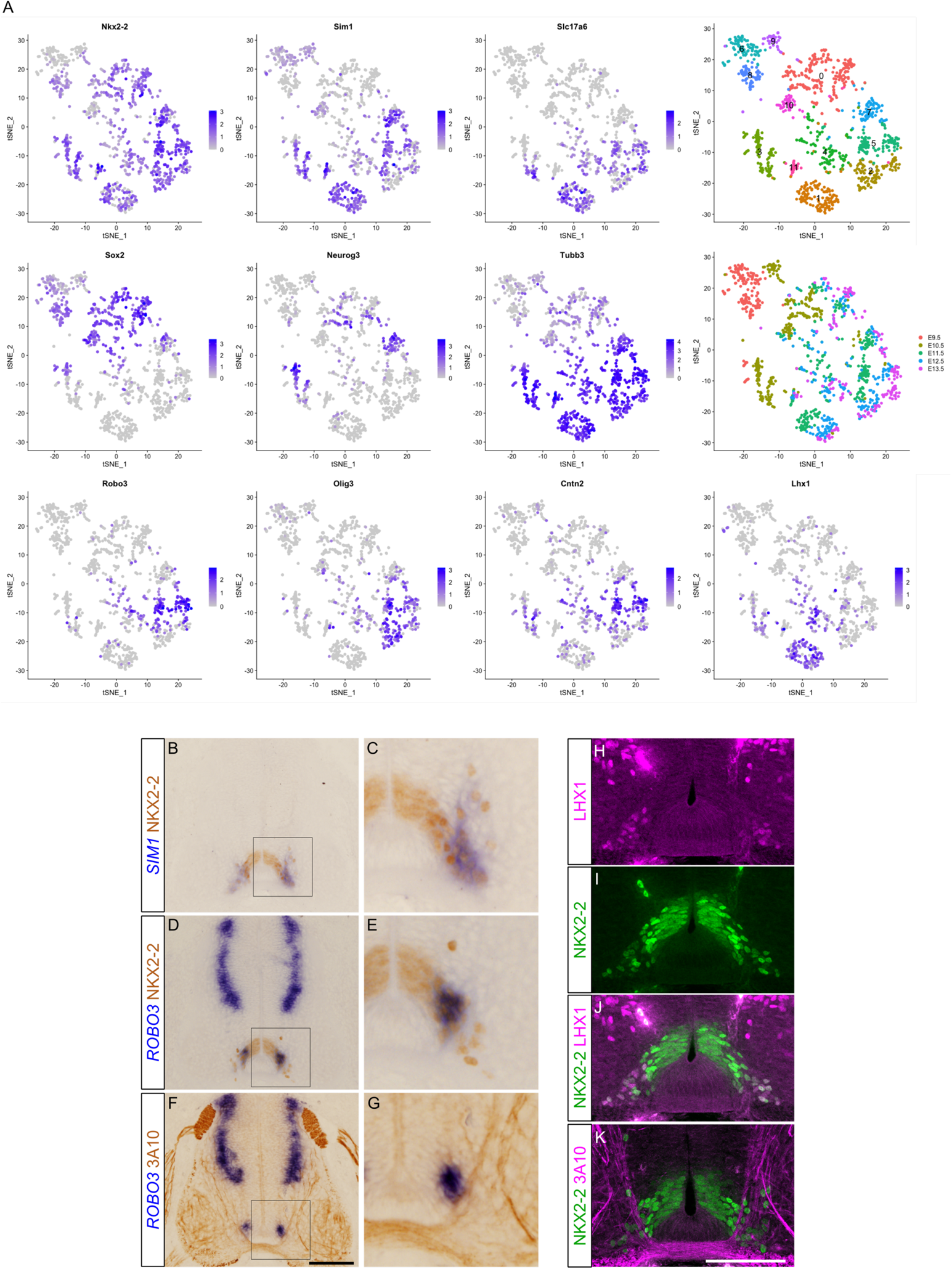
Two distinct subtypes of V3 INs in amniotes. (A) tSNE plot showing the cells with V3 IN identity derived from mouse embryonic spinal cord. Expression levels of genes indicated are visualized on the tSNE plot. Top right panel shows the result of graph-based clustering with each cluster being colored differently. Cluster numbers (0–11) are labeled. Middle right panel shows embryonic day when cells are corrected. (B–G) Expressions of NKX2-2, *SIM1* and *ROBO3* were examined in the chick spinal cord at the forelimb level at HH25–26. Growing axons were visualized by monoclonal antibody 3A10. Staining of *in situ* hybridization and immunohistochemistry was colored by blue and brown, respectively. C, E and G show the enlarged views of the boxed areas in B, D, and F, respectively. (H–J) Immunohistochemistry using NKX2-2 and LHX1 antibodies. (K) Immunohistochemistry using NKX2-2 and 3A10 antibodies. Ventral-most region of the spinal cord is shown in H–K. Scale bar: 100 µm in F for B, D and F, in K for H–K.

To distinguish these two subtypes *in vivo*, markers of each subtype (*ROBO3* and LHX1) were examined in the spinal cord of the chick embryo at Hamburger-Hamilton (HH) stage 25–26 (Fig.1B–K). NKX2-2 was used as a marker of the p3 progenitor domain and V3 INs. Post-mitotic V3 INs were distinguished by the expression of *SIM1*. We found that the expression of *ROBO3* was localized medially within post-mitotic V3 INs. To confirm this further, commissural axons passing through and bisecting the V3 INs were utilized as landmarks. Some V3 INs were observed laterally to the commissural axons; however, these cells did not express *ROBO3* (Fig.1F, G, K). Rather, *ROBO3* was expressed selectively by V3 INs located medially to the commissural axons (Fig.1F, G). In contrast, LHX1 was expressed in the laterally located population of the V3 INs (Fig.1H–J). These observations confirmed that two subtypes of V3 INs found by scRNA-seq are spatially, and probably also functionally, distinct populations in the developing spinal cord.

### Gene expression profile of V3 IN in zebrafish is distinct from that in amniotes

Next, we analyzed the scRNA-seq data derived from zebrafish whole embryos at 1 and 2 day post fertilization (dpf) (Farnsworth et al., 2020) to determine whether V3 INs of zebrafish are equivalent to those of amniotes. We extracted the data with the spinal cord profiles (detailed procedure of the data subsetting is provided in Fig.S1). Similarly to the data analysis of the mouse spinal cord, clustering and dimensionality reduction were applied (Fig.2). In the tSNE plot, the progenitor cells, which mainly consist of 1 dpf cells, were identified by the expression of *sox3* and non-expression of *elavl3* (*HuC*) (Fig.2, Clusters 0, 3, 4, 7, 11, 15, and 17). Expression of the domain specific TFs, such as *pax6b*, *olig2* and *nkx2.2b*, were localized at the specific space in the tSNE plot, which is parallel to the expression domain *in vivo* along the dorsoventral axis of the neural tube (Fig.2, Fig.S2). These data corroborate the evolutionary conservation of the progenitor domain organization in the neural tube (Cheesman and Eisen, 2004; Cheesman et al., 2004; Gribble et al., 2007; Guner and Karlstrom, 2007; Lewis et al., 2005; Park et al., 2002; Schäfer et al., 2005). V3 INs can be distinguished by the expression of *sim1a* (magenta circle in Fig.2). These V3 INs were *slc17a6b* positive glutamatergic neurons, like amniotes. However, unlike amniotes, almost all V3 INs expressed *robo3* in zebrafish, and there was no expression of *olig3* and *cntn2*. The expression of *Lhx1* and *Robo3* was mutually exclusive in the amniotes, yet this was not the case in the zebrafish. Furthermore, *neurog3*, one of the V3 IN specific TFs (Carcagno et al., 2014), was not expressed in the V3 INs of zebrafish (Fig.2, compared to Fig.1). These results indicate that V3 INs of zebrafish are not equivalent to those of amniotes so far as the gene expressions are compared.

**Figure 2.**
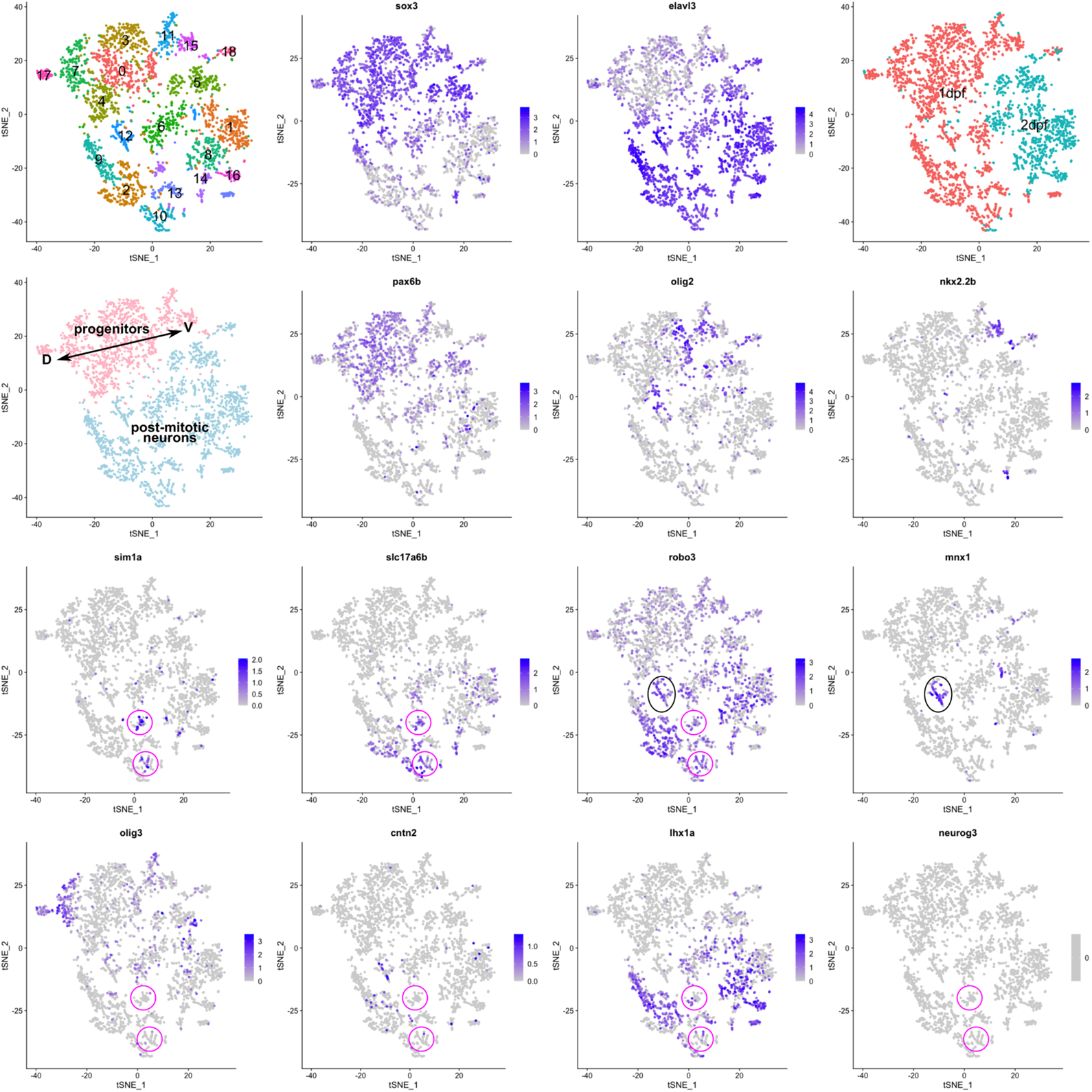
Gene expression profile of V3 IN in zebrafish is distinct from that in amniotes. (A) tSNE plot showing the cells with spinal cord identity derived from zebrafish embryos (1 and 2 dpf). Top left panel shows the result of graph-based clustering with each cluster being colored differently. Cluster numbers (0–18) are labeled. Top right panel shows embryonic day when cells are corrected. Left-most panel in second row shows progenitors and post-mitotic neurons by distinct color. D and V indicate dorsal and ventral, respectively. Other panels show the expression levels of genes indicated. Magenta and black circles indicate V3 INs and MNs, respectively.

### CRM in Robo3 locus is not conserved in teleosts

Focusing on *robo3* in zebrafish, *mnx1* positive MNs (black circle in Fig.2) also expressed *robo3*. This is clearly different from amniotes, in which *Robo3* is never expressed in MNs (Friocourt et al., 2019), implying that the transcriptional regulation of *Robo3* has diverged during vertebrate evolution. To confirm this, we searched for the transcription regulatory elements around *Robo3* locus using the available ChIP-seq and ATAC-seq data of neural cells (Fig.3). We found the putative *cis*-regulatory module (CRM) harboring multiple TF binding sites in an open chromatin state at around 20 kb upstream of *Robo3* transcription start site (TSS) (Fig. 3, CRM in *Robo3* locus is abbreviated as *Robo3-CRM*). To confirm the function of *Robo3-CRM*, reporter assay was performed using chick *in ovo* electroporation system. We cloned *Robo3-CRM* from mouse genome into vector containing minimal promoter and *GFP* (resulting construct is named *Robo3-CRM::GFP*). In the chick spinal cord after the electroporation of *Robo3-CRM::GFP*, GFP expression was observed to overlap the endogenous *ROBO3* expression, and GFP positive axons crossed the midline (Fig.3). More specifically, we found that GFP was expressed in LHX1 positive dorsal and intermediate INs, EVX1 positive V0 INs, and NKX2-2 positive V3 INs, but not in ISL1/2 positive dI3 INs or MNs, consistent with the endogenous *ROBO3* expressing cells (Fig.3) (Tulloch et al., 2019). These results demonstrate that *Robo3-CRM* can recapitulate the normal expression pattern of *Robo3* almost completely in the spinal cord.

**Figure 3.**
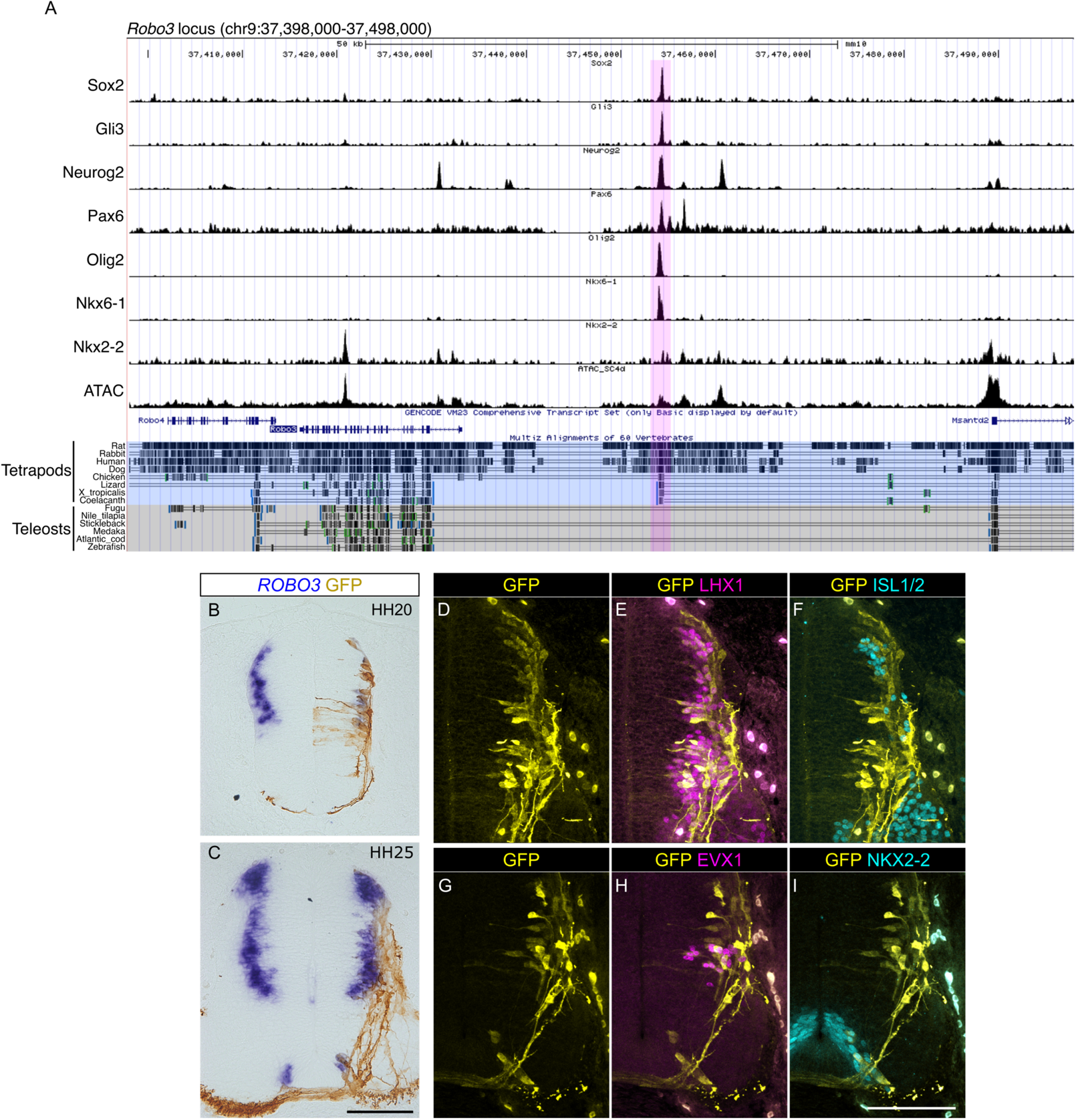
CRM in *Robo3* locus is not conserved in teleosts. (A) ChIP-seq and ATAC-seq peak call results are displayed in UCSC genome browser with Multiz Alignments track. *Robo3* locus in mouse genome (mm10) is displayed. Region harboring multiple TF binding sites (*Robo3-CRM*) is highlighted by magenta. Maltiz Alignment track of tatrapods and teleosts are highlighted by different colors. (B–I) *Robo3-CRM::GFP* was electroporated into chick neural tube. (B,C) Expression of GFP and *ROBO3* was examined at HH20 (B) and HH25 (C) after electroporation. (D–I) Expression of GFP and TFs indicated was examined at HH25 after electroporation. Scale bar: 100 µm in C for B and C, in I for D–I.

Notably, when sequence conservation was checked using Multiz alignments in UCSC genome browser among vertebrates (Blanchette et al., 2004), *Robo3-CRM* was conserved in tetrapods, but not in teleosts (Fig.3). Weak conservation of *Robo3-CRM* was observed in spotted gar, a basal actinopterygian (Fig.S3), suggesting that common ancestor of bony vertebrates possess *Robo3-CRM*, which has been lost in teleost lineage (Lee et al., 2011). These results suggest that divergent expression patterns of *robo3* in zebrafish are likely due to loss of *Robo3-CRM*. In addition, divergence of *Robo3-CRM* suggests that a part of axon guidance mechanisms in the post-mitotic neurons might have diverged during vertebrate evolution.

### Non-conservation of GBSs in Gli1 and Ptch1 loci in vertebrates

Next, we examined whether the upstream process before the progenitor fate specification has also diverged among vertebrates. Previous studies have reported that the impacts of loss of Gli function on the neuronal progenitors are different between mice and zebrafish (Chandrasekhar et al., 1999; Karlstrom et al., 2003; Tyurina et al., 2005; Vanderlaan et al., 2005; Wang et al., 2013). This raises the possibility that some Gli binding sites (GBSs), that is, the sites of action of Hh signaling, vary among vertebrates. To test this, *Gli1* and *Ptch1* loci were interrogated, as these genes are direct targets of Hh signaling. We searched for GBS in mouse genome and checked the sequence conservation using Maltiz alignments in the UCSC genome browser. Using Gli1 and Gli3 ChIP-seq data, many GBSs were identified in mouse *Gli1* and *Ptch1* loci near TSS and within intron (Fig.4), confirming previous reports (Dai et al., 1999; Nishi et al., 2015). ATAC-seq data confirmed that these GBSs are in an open chromatin state in neural cells. We found that most of these GBSs are not conserved among vertebrates. In the case of *Gli1*, clustered GBSs are conserved only in mammals, and in the *Ptch1* locus, most GBSs are conserved only in tetrapods, but were lost in teleosts (Fig.4, highlighted region). These findings imply that GBS is highly flexible element. This may partly explain the discrepancy of the *Gli* loss of function phenotypes between mice and zebrafish.

**Figure 4.**
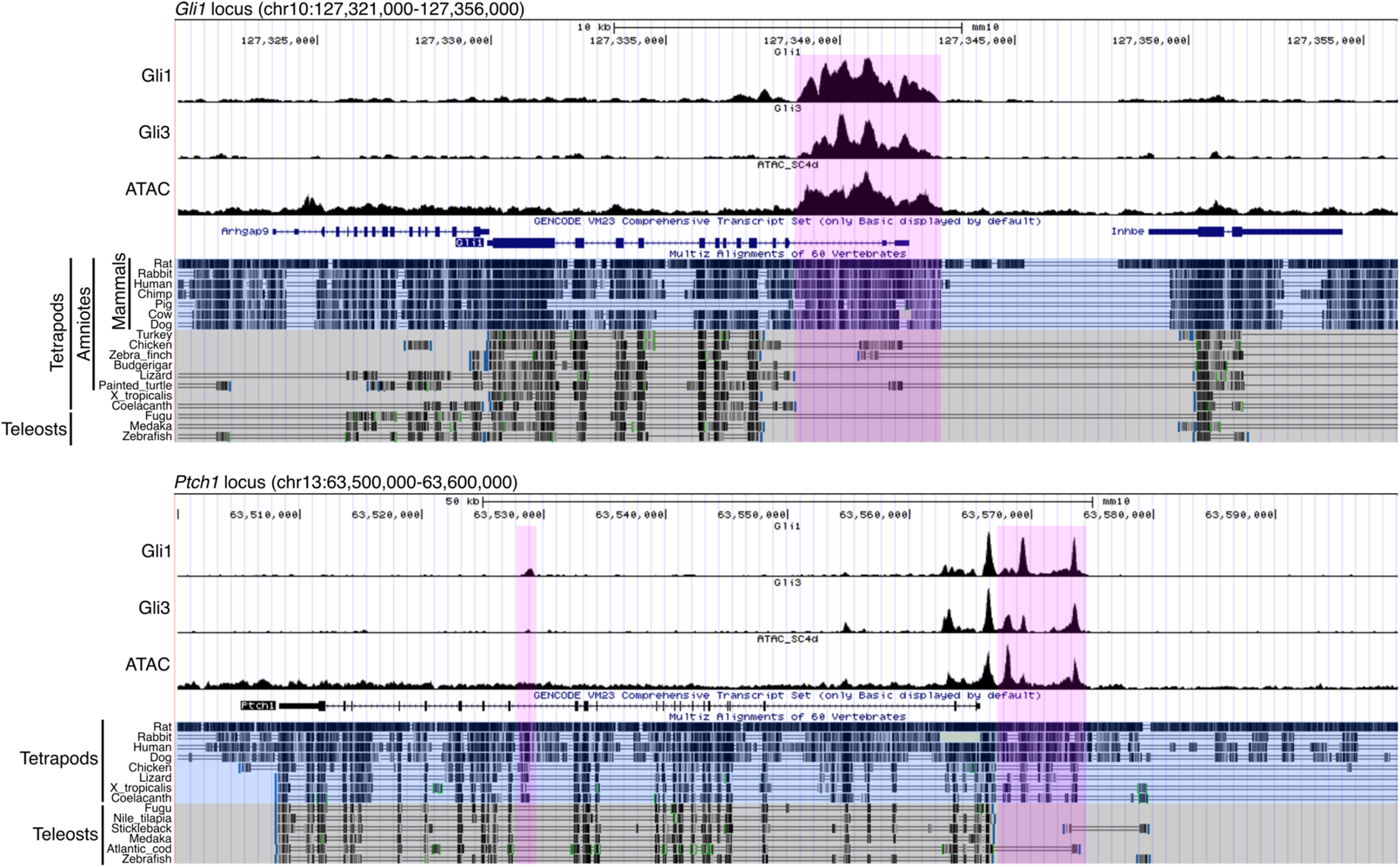
Non-conservation of GBSs in *Gli1* and *Ptch1* loci in vertebrates. Gli1/3 ChIP-seq and ATAC-seq peak call results are displayed in UCSC genome browser with Multiz Alignments track. *Gli1* and *Ptch1* loci in mouse genome (mm10) are shown. In *Gli1* locus, magenta region indicates the open chromatin region which is bound by Gli1/3, and is conserved only in mammals. Maltiz Alignment track of mammals and non-mammals are highlighted by different colors. In *Ptch1* locus, magenta regions indicate the open chromatin regions which are bound by Gli1/3, and are conserved only in tetrapods. Maltiz Alignment track of tatrapods and teleosts are highlighted by different colors.

### Non-conservation of CRMs of progenitor fate specifying TFs in vertebrates

Since the regulatory elements in *Robo3*, *Gli1* and *Ptch1* are not conserved among vertebrates, we expect that non-conservation of the regulatory elements can be found in other genes. Thus, we applied the same analysis to TFs functioning as progenitor fate determinants expressed in the neural tube. Indeed, we found that *Pax6-CRM*, *Gsx1-CRM*, *Dbx2-CRM*, *Irx3-CRMs* (2 CRMs, designated *Irx3-CRM1* and *2*), and *Olig2-CRMs* (2 CRMs, designated *Olig2-CRM1* and *2*) are conserved only in tetrapods, but were lost in teleosts (Fig.5, highlighted regions). These CRMs are in an open chromatin state in most cases, and are bound by Gli1, Gli3, Sox2, Neurog2, Pax6, Pax7, Olig2, Nkx2-2 and Nkx6-1 in various combinations (Fig.5). These TFs are essential components of GRN, regulating the progenitor fate in the neural tube (Balaskas et al., 2012; Delás and Briscoe, 2020; Exelby et al., 2021; Kutejova et al., 2016). Thus, we consider that non-conservation of these CRMs and conservation of progenitor organization has necessarily entailed the CRM turnover and GRN rewiring during vertebrate evolution. To exclude the possibility that these CRMs has been lost due to having no functions, we carried out the reporter assay by means of the chick *in ovo* electroporation system. This experiment confirmed that *Pax6-CRM*, *Gsx1-CRM*, *Dbx2-CRM*, *Irx3-CRM2*, and *Olig2-CRM2* function as enhancers in the neural tube cells, although more regulatory elements are needed to precisely recapitulate the endogenous expression pattern (Fig.6). Note that this method cannot capture the microRNA-based translational repression, which indeed affects the dorsal boundary of Olig2 (Chen et al., 2011). The only exception was *Irx-CRM1*, which exerted no enhancer function in the chick neural tube (data not shown). Enhancer function of *Olig2-CRM1* was already validated previously (Exelby et al., 2021; Oosterveen et al., 2012; Peterson et al., 2012; Wang et al., 2011). These results indicate that these CRMs are functional in amniotes, but nevertheless were lost in teleosts. This supports the notion that the progenitor fate specification system in the neural tube has undergone CRM turnover and GRN rewiring during vertebrate evolution (Fig.7).

**Figure 5.**
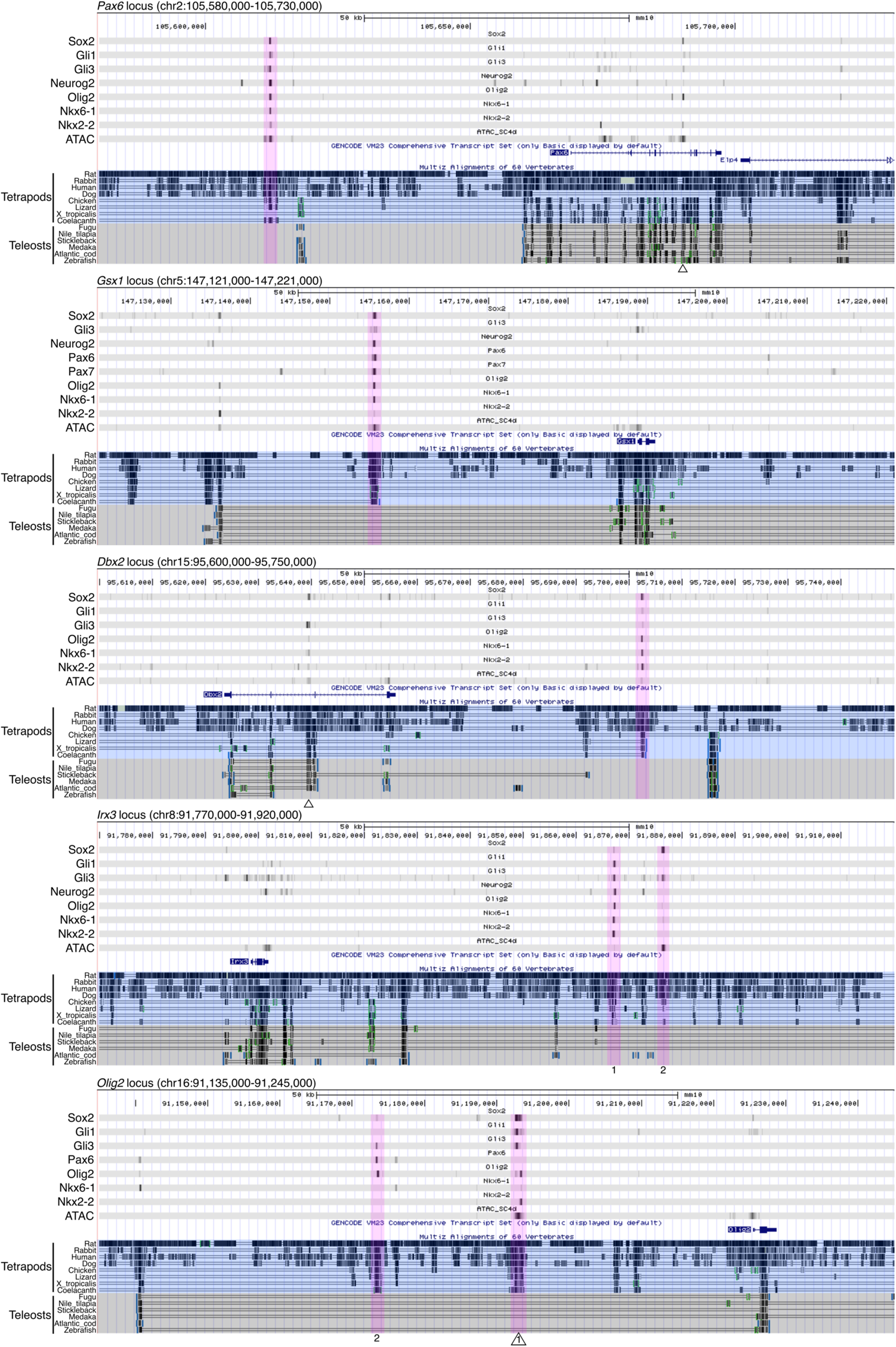
Non-conservation of CRMs of progenitor fate specifying TFs in vertebrates. ChIP-seq and ATAC-seq peak call results are displayed in UCSC genome browser with Multiz Alignments track (display mode is dense). Mouse *Pax6*, *Gsx1*, *Dbx2*, *Irx3* and *Olig2* loci are displayed. Regions harboring multiple TF binding sites (CRMs) are highlighted by magenta, and are conserved only in tetrapods. In *Irx3* and *Olig2* loci, two distinct CRMs are numbered 1 and 2 under the track. In *Pax6*, *Dbx2* and *Olig2* loci, previously validated CRMs are indicated by open triangle under the track (Oosterveen et al., 2012; Peterson et al., 2012).

**Figure 6.**
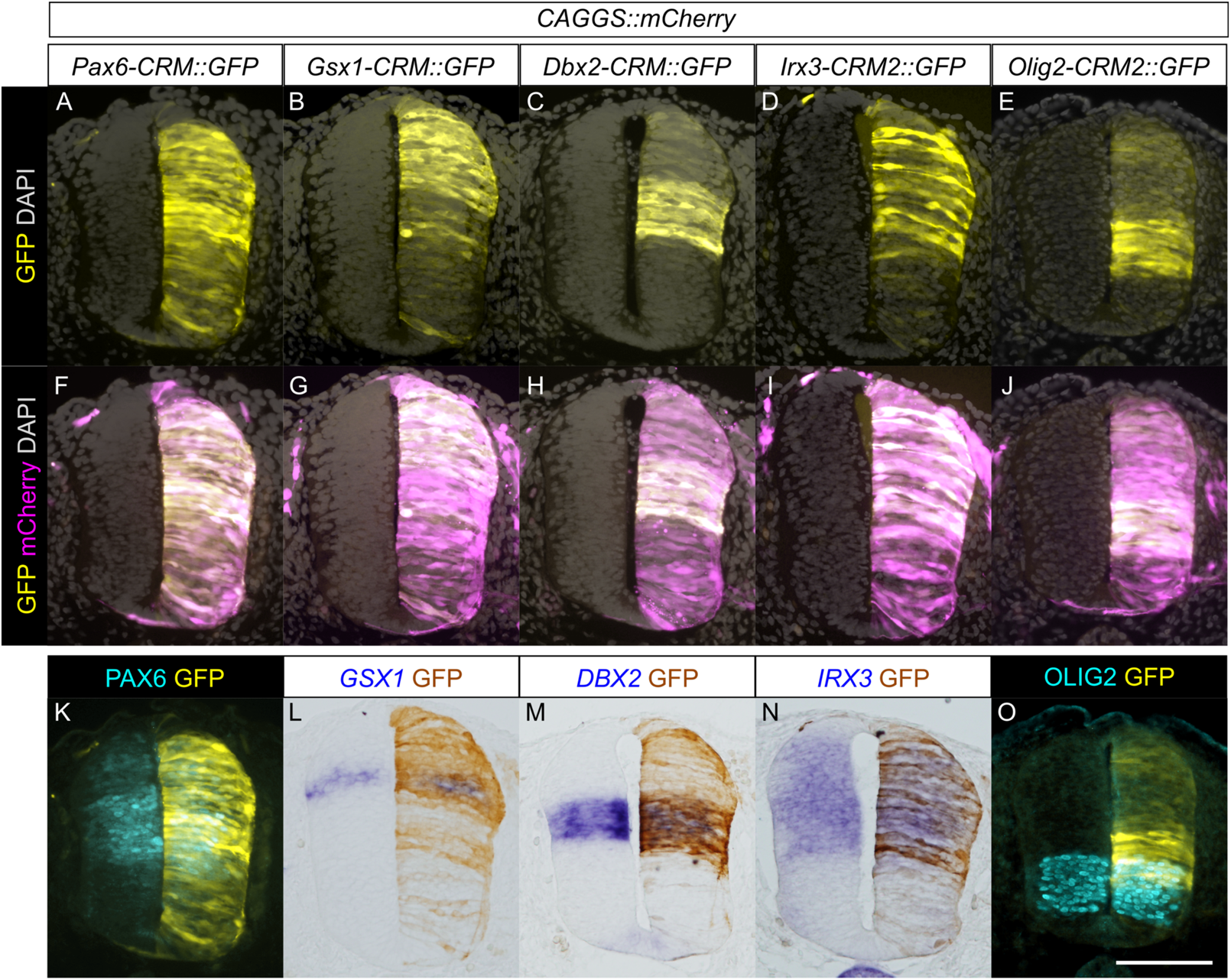
Enhancer functions of CRMs of progenitor fate specifying TFs. CRM reporter vectors indicated were electroporated into chick neural tube together with *CAGGS::mCherry* as control vector. (A–J) GFP and mCherry expression were examined at HH19-20 in the forelimb level neural tube. (K–O) The endogenous expression of genes indicated were examined by *in situ* hybridization (L, M, N) or immunohistochemistry (K, O) together with GFP expression. These five CRMs displayed enhancer functions, though GFP expression domains incompletely overlapped with endogenous expression domain. *Pax6-CRM* induced GFP expression almost ubiquitously, yet the expression was not observed in the roof plate (A, F, K). *Gsx1-CRM* induced GFP expression in the dorsal neural tube, but did not induce in the more ventral region than the endogenous expression domain (B, G, L). GFP expression domains induced by *Dbx2-CRM* or *Irx3-CRM2* overlapped with the endogenous expression domain almost completely (C, D, H, I, M, N). *Olig2-CRM2* induced the GFP expression in the intermediate to ventral region, which partially overlapped with, but was more dorsal than, the endogenous expression domain (E, J, O). Scale bar: 100 µm.

**Figure 7.**
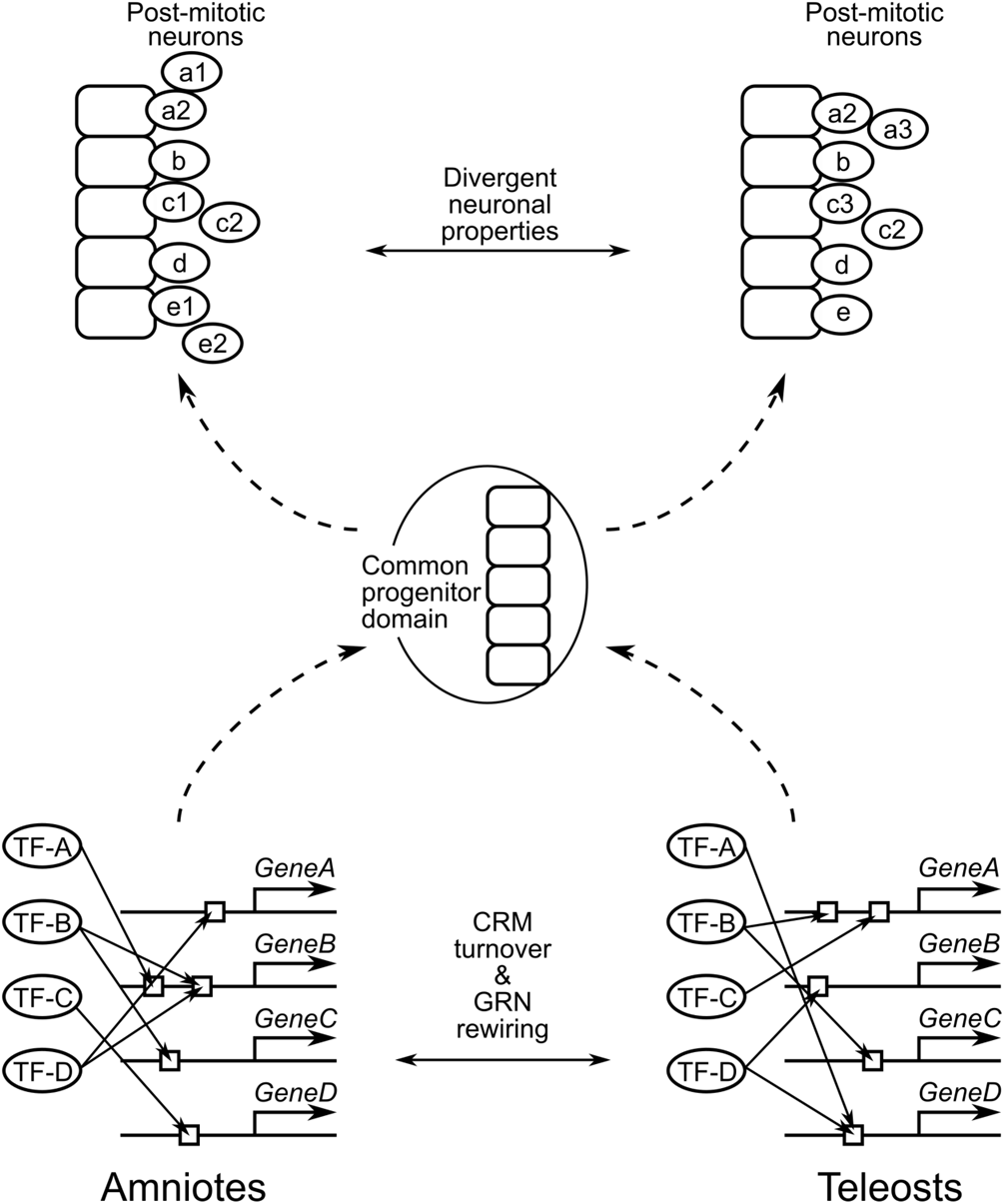
The hourglass-like pattern of the developmental divergence of the spinal cord. A summary of the present study is shown. In this drawing, development proceeds from bottom to top. Only five progenitor domains are set up, for simplification. Bottom drawing represents the GRNs regulating the progenitor domain establishment. CRMs (indicated by open boxes) located in the progenitor fate specifying genes (*GeneA–D*) has diverged (turnover) between amniotes and teleosts. Accordingly, these GRNs have been rewired, which is considered to be DSD. Nevertheless, these distinct GRNs result in the same progenitor domain organization due to the developmental system robustness. Top drawing represents the divergence of the differentiation process of post-mitotic neurons. After individual cells leave the progenitor domains as post-mitotic neurons, some neurons undergo distinct maturation process between amniotes and teleosts. Thus, there exist neuronal subpopulations whose function is different between amniotes and teleosts.

## Discussion

### Divergent properties of post-mitotic neurons in vertebrates

First, in the current study, we compared the single-cell transcriptomes of mice and zebrafish to elucidate the divergence of the post-mitotic neuronal differentiation. We found that, in amniotes, V3 INs can be divided into two distinct subtypes. One is located medially and expresses *Robo3*, *Olig3*, and *Cntn2*, and the other is located laterally and expresses *Lhx1* (Fig.1). We tried to make these V3 INs correspond with those of zebrafish, but were unsuccessful (Fig.1,2). This indicates that V3 INs of amniotes is not equivalent to those of zebrafish, at least regarding gene expression profile, which supports the hypothesis that the properties of post-mitotic neurons in the spinal cord have diverged among vertebrates. This is in contrast to the case of progenitor cells, in which each progenitor domain of zebrafish readily corresponds with those of amniotes (Fig.S2).

The role of V3 INs in mice is to secure the stable locomotor rhythm (Zhang et al., 2008). Several studies have also suggested that V3 INs contribute to left-right synchronous motor output, such as gallop and bound (Danner et al., 2019; Kiehn, 2016; Rabe et al., 2009). These gaits can be expressed by quadrupeds, but not by fish. This suggests that the different locomotor behavior between amniotes and teleosts is associated with the divergence of V3 IN properties. The same may be the case with V2a INs, whose roles have diverged between amniotes and teleosts (Azim et al., 2014; Crone et al., 2008, 2009; Eklöf-Ljunggren et al., 2012; Kiehn, 2016). Given that the numbers of muscles and corresponding MNs have significantly increased during tetrapod evolution, especially regarding limb muscles (Diogo and Abdala, 2010), additional layers of neuronal circuits were required to precisely control complex locomotor behavior (Kiehn, 2016). This may be accomplished, at least in part, by tinkering with the already existing neuronal population (Jacob, 1977), eventually leading to the divergence of the post-mitotic neuronal properties among vertebrates. Distinct expression profiles of *Robo3* between amniotes and zebrafish may be one of the strategies for neuronal tinkering (Fig.1–3).

### Divergent processes leading to the conserved progenitor domains

In the latter part of this study, we investigated the transcriptional regulatory elements located around the progenitor fate specifying genes in order to reveal to what extent the upstream process before the progenitor specification has diverged in vertebrates. In *Gli1* locus, clustered GBSs near the TSS are conserved only in mammals, but not in teleosts (Fig.4). Even in birds and reptiles, these GBSs are conserved only partially. Nevertheless, *Gli1* is expressed in response to Hh signaling commonly in vertebrates (Aglyamova and Agarwala, 2007; Karlstrom et al., 2003; Luo et al., 2006), suggesting the presence of distinct GBSs with equivalent function. Indeed, zebrafish possesses three GBSs in *Gli1* locus, which are conserved only in some teleost species (Wang et al., 2013), indicating that regions where Hh signaling is eventually transduced has shifted during vertebrate evolution. This is supported by a similar situation in *Ptch1* locus (Fig.4) (Wang et al., 2013). Bicoid is a morphogen that plays a crucial role for the patterning of the anterior-posterior axis in *Drosophila* blastoderm, and there are fundamental similarities between Bicoid and Shh (Briscoe and Small, 2015). It is noteworthy that Bicoid binding sites have also undergone rapid turnover in Diptera (McGregor et al., 2001). The flexibility of morphogen response elements might contribute to the integration of the morphogen dependency into the patterning system of the embryo (Dearden and Akam, 1999; Miyamoto and Wada, 2013; Ren et al., 2020; Stauber et al., 1999).

In *Pax6*, *Gsx1*, *Dbx2*, *Irx3*, and *Olig2* loci in mouse genome, we identified functional CRMs bound by multiple TFs (Fig.5,6). These CRMs are conserved only in tetrapods, but were lost in teleosts (Fig.5). Given that the progenitor domain organization is conserved among vertebrates, these findings indicate the CRM turnover and GRN rewiring during vertebrate evolution (Fig.7). This is considered to be a case of developmental system drift (DSD) (True and Haag, 2001). Several studies have reported similar situations, in which divergent regulatory sequences in different species result in the conserved gene expressions (Barrière et al., 2012; Fisher et al., 2006; Hare et al., 2008; Ludwig et al., 1998, 2000; Paris et al., 2013; Stolfi et al., 2014; Swanson et al., 2011). In these cases, including the present study, GRN architecture is likely to be maintained as a whole despite the CRM turnover, as demonstrated in mammalian evolution (Stergachis et al., 2014; Vierstra et al., 2014).

The process of the neural tube formation is different between amniotes and zebrafish not only genetically, but also morphogenetically. In amniotes, a midline groove is formed by bending of the neural plate, the edges of which are then fused, thus forming the neural tube (Schoenwolf and Smith, 1990). On the other hand, in zebrafish, the solid neural keel with no central canal is formed by convergent movement of the neural plate cells, and then the central canal opens secondarily (Schmitz et al., 1993). Taken together, the neural tube formation is a case of DSD from both genetic and morphogenetic viewpoints.

### Molecular basis of the hourglass model in the spinal cord

Taken together, our results demonstrate that divergence of the developmental process of the spinal cord is in accordance with the developmental hourglass model. Then, what impose the hourglass-like pattern on the evolution of spinal cord development? The progenitor regionalization in the neural tube is a highly robust developmental system, and thus is insensitive against noises and mutations (Balaskas et al., 2012; Delás and Briscoe, 2020; Exelby et al., 2021; Xiong et al., 2013; Zagorski et al., 2017). Simulation studies have also proposed that the developmental system robustness is an emergent property of the complex GRN (Bergman and Siegal, 2003; von Dassow et al., 2000; Siegal and Bergman, 2002), indirectly supporting the robustness of the neuronal progenitor specification system, which is organized by highly complex GRN (Kutejova et al., 2016). From the evolutionary perspective, such a robust (canalized) developmental system can accommodate genetic variations without disturbing development (Rutherford and Lindquist, 1998; Waddington, 1942, 1957). In other words, the progenitor specification system in the neural tube buffers the genetic variations. We speculate that this situation consequently led to the CRM turnover and GRN rewiring, while the progenitor arrangement was maintained. Once cells are released from the progenitor domains as post-mitotic neurons, individual neurons depend on the mechanisms regulating post-mitotic maturation which is distinct from the progenitor specification GRN. As mentioned before, the maturation process is considered to diverge in association with locomotor divergence. Eventually, through vertebrate evolution, developmental divergence of the spinal cord might lead to the hourglass shape. The focus of the current study was the spinal cord. Thus, it is of particular interest to examine whether this scenario is also the case in other developmental systems or even in whole embryonic level.

More importantly, if partial modification is admitted, neuroectodermal regionalization is conserved among bilaterians beyond vertebrates (Arendt, 2018; Denes et al., 2007; Jung and Dasen, 2015). This suggests that the developmental system robustness of neuroectodermal regionalization had already been acquired in the last common ancestor of bilaterians, and that upstream signals of the neuroectodermal regionalization had been modified after divergence of phyla (e.g., Hh signalling recruitment in deuterostomes) (Miyamoto and Wada, 2013; Ren et al., 2020). This is supported by the notion of the common origin of the central nervous system (CNS) in bilaterians (Arendt, 2018; Arendt et al., 2016; Denes et al., 2007), but is inconsistent with the convergent evolution of CNS (Martín-Durán et al., 2018).

## Acknowledgements

We thank Y. Watanabe and T. Shimada for technical supports. This work was supported by JSPS KAKENHI Grant Number 18K14718 and 18K06835. Computations were partially performed on the NIG supercomputer at ROIS National Institute of Genetics. The monoclonal antibodies 74.5A5, 4F2, 3A10, 39.4D5, 99.1-3A2, PAX6, and DSHB-GFP-4C9, developed by T. M. Jessell, J. Dodd, S. Brenner-Morton, A. Kawakami and DSHB were obtained from the Developmental Studies Hybridoma Bank, created by the NICHD of the NIH and maintained at The University of Iowa, Department of Biology, Iowa City, IA 52242.

## Author Contributions

Conceptualization, K.M.; Methodology, K.M.; Formal Analysis, K.M.; Investigation, and C.S.; Writing - Original Draft, K.M.; Writing - Review & Editing, K.M and H.Y.; Funding Acquisition, K.M. and H.Y.

## Methods

### Experimental animals

Fertilized chicken eggs were obtained from Takeuchi farm (Nara, Japan) and incubated at 38°C in a humidified incubator. Embryos were staged according to Hamburger-Hamilton stage series (Hamburger and Hamilton, 1951). All animal experiments were performed in accordance with the Rules of Fukushima Medical University Animal Experiments, with approval of the Animal Experiments Committee of Fukushima Medical University (approval number 2020103).

### scRNA-seq data analysis

Mouse spinal cord scRNA-seq data (Delile et al., 2019) was obtained from ArrayExpress (accession number E-MTAB-7320). UMI count matrix was generated using Cell Ranger version 3.1.0 (10x Genomics). In this step, normalization was skipped in order to maximize sensitivity (cellranger aggr was executed with --normalize=none). The output matrix was fed into Seurat (R package) version 3.1.5 (Butler et al., 2018). Zebrafish whole embryo scRNA-seq data (Farnsworth et al., 2020) was obtained as Seurat object (.rds file) from Dr. Miller’s website (https://www.adammillerlab.com/resources-1). Detailed procedures of normalization, data subsetting, graph-based clustering, and dimensionality reduction using Seurat are provided in https://github.com/kmukaigasa/Spinalcord_MouseZebrafish.

### ChIP-seq data analysis

ChIP-seq data were obtained from NCBI sequence read archive (SRA). The accession numbers are as follows: GSE66961 for Pax6 (Sun et al., 2015); GSE42132 for Sox2 and Gli1 (Peterson et al., 2012); GSE61673 for Gli3, Nkx2-2, Nkx6-1, and Olig2 (Nishi et al., 2015); GSE114172 for Neurog2 (Aydin et al., 2019); and GSE87180 for Pax7 (Mayran et al., 2018). The cell types used were embryonic forebrain (Pax6), AtT-20 cell (Pax7), and neural cells differentiated from mouse ES cells (Sox2, Gli1/3, Nkx2-2, Nkx6-1, Olig2, Neurog2). Sequencing adaptors and low quality bases were trimmed by Trimmomatic (version 0.39). FastQC (version 0.11.9) was used for sequence quality check. Read mapping to mouse reference genome (mm10) was performed by Bowtie2 (version 2.3.5) with default setting. Reads with low mapping quality (MAPQ < 10) were removed by samtools (version 1.9). Peak call was performed by MACS2 (version 2.2.6). Output bedGraph file was converted to BigWig format by bdg2bw (https://gist.github.com/jl32587/34370c995460f9d5ad65). BigWig track was visualized in UCSC genome browser.

### ATAC-seq data analysis

ATAC-seq data (Metzis et al., 2018) were obtained from ArrayExpress (E-MTAB-6337). From the dataset, data of the neural progenitor cells with spinal cord identity were used. Read quality control was done as ChIP-seq. Read mapping was performed by Bowtie2 with the following parameters; -X 2000 --sensitive-local. PCR duplicates were marked using Picard (version 2.9.2). Reads with low mapping quality (MAPQ < 30) were removed by samtools. Peak call was performed by MACS2 with following parameters; -f BAMPE -g mm -q 0.05 --nomodel --keep-dup auto -B.

### Comparison of genomic sequence by VISTA

For interspecies comparison of genomic sequence, partial genomic sequences were obtained from UCSC genome browser (https://genome.ucsc.edu) and Ensembl (https://www.ensembl.org/index.html), and VISTA (Frazer et al., 2004) are used to align and visualize the results.

### Vector construction

For RNA probe preparation, partial fragments of *ROBO3*, *IRX3*, *DBX2*, and *GSX1* were amplified by PCR from chick embryonic spinal cord cDNA. The primer sequences used are provided in Table S2. The fragments were inserted into pCR-XL-TOPO (Thermo Fisher) or *pBluescriptKS*. For reporter assay of CRMs, genomic DNA fragments were amplified by PCR from a solution which was prepared by digesting the tail tip of a C57BL/6J mouse. The amplified fragments were inserted into *pSF-pA-MinProm-eGFP* vector (Oxford genetics). Positions of CRMs within the mouse reference genome are provided in Table S3. For construction of *pCAGGS-mCherry*, *mCherry* gene cassette of *pmCherry-C1* (Clontech) was inserted into *pCAGGS* (Niwa et al., 1991).

### in ovo electroporation

A small window was opened on the top of the fertilized chicken egg shell. The electrodes were placed on both sides of the neural tube of a HH12-13 chick embryo. Plasmid DNA was injected into the neural tube. During injection, electric pulses (25 V, 50 ms, five times, 950 ms interval) were applied using CUY21EDIT (BEX, Tokyo, Japan). Plasmids were prepared at 1 µg/µl for *pSF-Robo3-CRM::GFP*, 20–50 ng/µl for *pSF-Pax6-CRM::GFP*, *pSF-Gsx1-CRM::GFP*, *pSF-Dbx2-CRM::GFP*, *pSF-Irx3-CRM::GFP*, *pSF-Olig2-CRM::GFP*, 0.5 ng/µl for *pCAGGS-mCherry*.

### Immunohistochemistry

Embryos were fixed in 0.1 M phosphate buffer / 4% paraformaldehyde (PFA) at room temperature for 45–60 min or 4°C overnight. The fixed embryos were then cryoprotected in 20% sucrose at 4°C overnight, embedded in OCT compound, and cryosectioned. In cases of overnight fixed samples, sections were boiled for 20 min in sodium citrate buffer (10 mM sodium citrate, 0.05% Tween 20, pH 6.0) and cooled down to room temperature. After washing with phosphate-buffered saline (PBS) containing 0.1% Triton X-100 (PBST), the sections were incubated with primary antibodies at 4°C overnight, washed with PBST three times for 5 min, and incubated with secondary antibodies for 1–2 h at room temperature. After washing with PBST three times for 5 min, slides were coverslipped in VECTASHIELD (Vector laboratories). The antibodies used are described in Table S4. In the cases of anti-Pax6 and anti-Lhx1, Can Get Signal (TOYOBO, NKB-401) was used for the diluent instead of PBST. Images were captured using an Olympus BX51 fluorescent microscope equipped with an Olympus DP71 digital camera, or an Olympus FluoView FV1000 confocal microscope.

### in situ hybridization

For double staining of immunohistochemistry and *in situ* hybridization, immunohistochemistry was performed as described above with primary and secondary antibody incubation for 1 h at room temperature, and signal was developed by VECTASTAIN Elite ABC kit (Vector laboratories). Following that, the process of *in situ* hybridization, as described below, was started. Plasmid for RNA probe template was linearized, and DIG-labeled RNA probes were generated using DIG RNA labeling mix (Sigma-Aldrich, 11277073910) and T3 RNA polymerase (Promega, P2083).

Tissue slides were washed with PBS for 5 min, treated with protainaseK (2µg/ml) for 10 min, washed with PBS for 5 min, fixed in 4% PFA for 10 min, and washed three times with PBS for 5 min. The slides were then incubated in an acetylation solution (100 mM triethanolamine, pH 8.0) twice for 2–3 min, transferred in new acetylation solution and acetylated for 15 min by adding dropwise acetic anhydride (0.3% final concentration), and washed with PBS three times for 5 min. The slides were prehybridized with hybridization buffer (50% formamide, 5X SSC, 5X Denhardt’s, 500 µg/ml herring sperm DNA, 250 µg/ml yeast RNA) for 2–3 h at room temperature, and hybridized with hybridization buffer containing DIG-labelled RNA probe. The slides were coverslipped and incubated in a humidified chamber at 70°C overnight. The following day, the slides were transferred in 5X SSC at 70°C for 5 min, washed twice in 0.2X SSC for 30 min at 70°C, and washed in 0.2X SSC for 30 min at room temperature. Slides were washed with buffer B1 (100mM Tris-HCl pH 7.5, 150 mM NaCl, 0.1% Tween 20) three times for 30 min, incubated with buffer B1 containing 10% heat-inactivated normal goat serum for 2–3 h at room temperature, and then incubated with buffer B1 containing alkaline phosphatase conjugated anti-DIG antibody (Roche 11093274910, 1:5000) and 1% heat-inactivated normal goat serum at 4°C overnight. The following day, slides were washed five times with buffer B1 for 30 min, and incubated in buffer B3 (100 mM Tris-HCl pH 9.5, 100 mM NaCl, 50 mM MgCl2, 0.1% Tween20) three times for 5 min. The slides were incubated with BM purple (Roche, 11442074001) until the signal was visualized (3 h to overnight). After that, the slides were washed three times with buffer B1 for 30 min, then washed three times with PBS for 5 min, and coverslipped in Fluoromount (Diagnostic Biosystems, K024).

## References

Aglyamova, G. V., and Agarwala, S. (2007). Gene expression analysis of the Hedgehog signaling cascade in the chick midbrain and spinal cord. Dev. Dyn. 236, 1363–1373.

Andrews, M.G., Kong, J., Novitch, B.G., and Butler, S.J. (2019). New perspectives on the mechanisms establishing the dorsal-ventral axis of the spinal cord. Curr. Top. Dev. Biol. 132, 417–450.

Arendt, D. (2018). Animal Evolution: Convergent Nerve Cords? Curr. Biol. 28, R225–R227.

Arendt, D., Tosches, M.A., and Marlow, H. (2016). From nerve net to nerve ring, nerve cord and brain-evolution of the nervous system. Nat. Rev. Neurosci. 17, 61–72.

Aydin, B., Kakumanu, A., Rossillo, M., Moreno-Estellés, M., Garipler, G., Ringstad, N., Flames, N., Mahony, S., and Mazzoni, E.O. (2019). Proneural factors Ascl1 and Neurog2 contribute to neuronal subtype identities by establishing distinct chromatin landscapes. Nat. Neurosci. 22, 897–908.

Azim, E., Jiang, J., Alstermark, B., and Jessell, T.M. (2014). Skilled reaching relies on a V2a propriospinal internal copy circuit. Nature 508, 357–363.

Balaskas, N., Ribeiro, A., Panovska, J., Dessaud, E., Sasai, N., Page, K.M., Briscoe, J., and Ribes, V. (2012). Gene regulatory logic for reading the Sonic Hedgehog signaling gradient in the vertebrate neural tube. Cell 148, 273–284.

Barrière, A., Gordon, K.L., and Ruvinsky, I. (2012). Coevolution within and between Regulatory Loci Can Preserve Promoter Function Despite Evolutionary Rate Acceleration. PLoS Genet. 8, e1002961.

Bergman, A., and Siegal, M.L. (2003). Evolutionary capacitance as a general feature of complex gene networks. Nature 424, 549–552.

Blanchette, M., Kent, W.J., Riemer, C., Elnitski, L., Smit, A.F.A., Roskin, K.M., Baertsch, R., Rosenbloom, K., Clawson, H., Green, E.D., et al. (2004). Aligning multiple genomic sequences with the threaded blockset aligner. Genome Res. 14, 708–715.

Briscoe, J., and Small, S. (2015). Morphogen rules: design principles of gradient-mediated embryo patterning. Development 142, 3996–4009.

Briscoe, J., and Thérond, P.P. (2013). The mechanisms of Hedgehog signalling and its roles in development and disease. Nat. Rev. Mol. Cell Biol. 14, 416–429.

Briscoe, J., Sussel, L., Serup, P., Hartigan-O’Connor, D., Jessell, T.M., Rubenstein, J.L.R., and Ericson, J. (1999). Homeobox gene Nkx2.2 and specification of neuronal identity by graded Sonic hedgehog signalling. Nature 398, 622–627.

Briscoe, J., Pierani, A., Jessell, T., and Ericson, J. (2000). A homeodomain protein code specifies progenitor cell identity and neuronal fate in the ventral neural tube. Cell 101, 435– 445.

Butler, A., Hoffman, P., Smibert, P., Papalexi, E., and Satija, R. (2018). Integrating single-cell transcriptomic data across different conditions, technologies, and species. Nat. Biotechnol. 36, 411–420.

Carcagno, A.L., Di Bella, D.J., Goulding, M., Guillemot, F., and Lanuza, G.M. (2014). Neurogenin3 Restricts Serotonergic Neuron Differentiation to the Hindbrain. J. Neurosci. 34, 15223–15233.

Challa, A.K., McWhorter, M.L., Wang, C., Seeger, M.A., and Beattie, C.E. (2005). Robo3 isoforms have distinct roles during zebrafish development. Mech. Dev. 122, 1073–1086.

Chandrasekhar, A., Schauerte, H.E., Haffter, P., and Kuwada, J.Y. (1999). The zebrafish detour gene is essential for cranial but not spinal motor neuron induction. Development 126, 2727–2737.

Cheesman, S.E., and Eisen, J.S. (2004). Gsh1 Demarcates Hypothalamus and Intermediate Spinal Cord in Zebrafish. Gene Expr. Patterns 5, 107–112.

Cheesman, S.E., Layden, M.J., Von Ohlen, T., Doe, C.Q., and Eisen, J.S. (2004). Zebrafish and fly Nkx6 proteins have similar CNS expression patterns and regulate motoneuron formation. Development 131, 5221–5232.

Chen, J.A., Huang, Y.P., Mazzoni, E.O., Tan, G.C., Zavadil, J., and Wichterle, H. (2011). Mir-17-3p Controls Spinal Neural Progenitor Patterning by Regulating Olig2/Irx3 Cross-Repressive Loop. Neuron 69, 721–735.

Crone, S.A., Quinlan, K.A., Zagoraiou, L., Droho, S., Restrepo, C.E., Lundfald, L., Endo, T., Setlak, J., Jessell, T.M., Kiehn, O., et al. (2008). Genetic Ablation of V2a Ipsilateral Interneurons Disrupts Left-Right Locomotor Coordination in Mammalian Spinal Cord. Neuron 60, 70–83.

Crone, S.A., Zhong, G., Harris-Warrick, R., and Sharma, K. (2009). In mice lacking V2a interneurons, gait depends on speed of locomotion. J. Neurosci. 29, 7098–7109.

Currie, P.D., and Ingham, P.W. (1996). Induction of a specific muscle cell type by a hedgehog-like protein in zebrafish. Nature 382, 452–455.

Dai, P., Akimaru, H., Tanaka, Y., Maekawa, T., Nakafuku, M., and Ishii, S. (1999). Sonic Hedgehog-induced Activation of the Gli1 Promoter Is Mediated by GLI3. J. Biol. Chem. 274, 8143–8152.

Danner, S.M., Zhang, H., Shevtsova, N.A., Borowska-Fielding, J., Deska-Gauthier, D., Rybak, I.A., and Zhang, Y. (2019). Spinal V3 Interneurons and Left–Right Coordination in Mammalian Locomotion. Front. Cell. Neurosci. 13, 516.

von Dassow, G., Meir, E., Munro, E.M., and Odell, G.M. (2000). The segment polarity network is a robust developmental module. Nature 406, 188–192.

Dearden, P., and Akam, M. (1999). Developmental evolution: Axial patterning in insects. Curr. Biol. 9, R591–R594.

Delás, M.J., and Briscoe, J. (2020). Repressive interactions in gene regulatory networks: When you have no other choice. Curr. Top. Dev. Biol. 139, 239–266.

Del Barrio, M.G., Taveira-Marques, R., Muroyama, Y., Yuk, D.-I., Li, S., Wines-Samuelson, M., Shen, J., Smith, H.K., Xiang, M., Rowitch, D., et al. (2007). A regulatory network involving Foxn4, Mash1 and delta-like 4/Notch1 generates V2a and V2b spinal interneurons from a common progenitor pool. Development 134, 3427–3436.

Delile, J., Rayon, T., Melchionda, M., Edwards, A., Briscoe, J., and Sagner, A. (2019). Single cell transcriptomics reveals spatial and temporal dynamics of gene expression in the developing mouse spinal cord. Development 146, dev173807.

Denes, A.S., Jékely, G., Steinmetz, P.R.H., Raible, F., Snyman, H., Prud’homme, B., Ferrier, D.E.K., Balavoine, G., and Arendt, D. (2007). Molecular Architecture of Annelid Nerve Cord Supports Common Origin of Nervous System Centralization in Bilateria. Cell 129, 277–288.

Ding, Q., Motoyama, J., Gasca, S., Mo, R., Sasaki, H., Rossant, J., and Hui, C.C. (1998). Diminished Sonic hedgehog signaling and lack of floor plate differentiation in Gli2 mutant mice. Development 125, 2533–2543.

Ding, Q., Joshi, P.S., Xie, Z.-H., Xiang, M., and Gan, L. (2012). BARHL2 transcription factor regulates the ipsilateral/contralateral subtype divergence in postmitotic dI1 neurons of the developing spinal cord. Proc. Natl. Acad. Sci. U. S. A. 109, 1566–1571.

Diogo, R., and Abdala, V. (2010). Muscles of vertebrates : comparative anatomy, evolution, homologies and development (CRC Press).

Duboule, D. (1994). Temporal colinearity and the phylotypic progression: a basis for the stability of a vertebrate Bauplan and the evolution of morphologies through heterochrony. Development 1994, 135–142.

Echelard, Y., Epstein, D.J., St-Jacques, B., Shen, L., Mohler, J., McMahon, J.A., and McMahon, A.P. (1993). Sonic hedgehog, a member of a family of putative signaling molecules, is implicated in the regulation of CNS polarity. Cell 75, 1417–1430.

Ekker, S.C., Ungar, A.R., Greenstein, P., von Kessler, D.P., Porter, J.A., Moon, R.T., and Beachy, P.A. (1995). Patterning activities of vertebrate hedgehog proteins in the developing eye and brain. Curr. Biol. 5, 944–955.

Eklöf-Ljunggren, E., Haupt, S., Ausborn, J., Dehnisch, I., Uhlén, P., Higashijima, S., and El Manira, A. (2012). Origin of excitation underlying locomotion in the spinal circuit of zebrafish. Proc. Natl. Acad. Sci. U. S. A. 109, 5511–5516.

Exelby, K., Herrera-Delgado, E., Garcia Perez, L., Perez-Carrasco, R., Sagner, A., Metzis, V., Sollich, P., and Briscoe, J. (2021). Precision of Tissue Patterning is Controlled by Dynamical Properties of Gene Regulatory Networks. Development 148, dev197566.

Farnsworth, D.R., Saunders, L.M., and Miller, A.C. (2020). A single-cell transcriptome atlas for zebrafish development. Dev. Biol. 459, 100–108.

Fisher, S., Grice, E.A., Vinton, R.M., Bessling, S.L., and McCallion, A.S. (2006). Conservation of RET regulatory function from human to zebrafish without sequence similarity. Science 312, 276–279.

Frazer, K.A., Pachter, L., Poliakov, A., Rubin, E.M., and Dubchak, I. (2004). VISTA: computational tools for comparative genomics. Nucleic Acids Res. 32, W273–W279.

Friocourt, F., and Chédotal, A. (2017). The Robo3 receptor, a key player in the development, evolution, and function of commissural systems. Dev. Neurobiol. 77, 876–890.

Friocourt, F., Kozulin, P., Belle, M., Suárez, R., Di-Poï, N., Richards, L.J., Giacobini, P., and Chédotal, A. (2019). Shared and differential features of Robo3 expression pattern in amniotes. J. Comp. Neurol. 527, 2009–2029.

Gribble, S.L., Nikolaus, O.B., and Dorsky, R.I. (2007). Regulation and function of Dbx genes in the zebrafish spinal cord. Dev. Dyn. 236, 3472–3483.

Guner, B., and Karlstrom, R.O. (2007). Cloning of zebrafish nkx6.2 and a comprehensive analysis of the conserved transcriptional response to Hedgehog/Gli signaling in the zebrafish neural tube. Gene Expr. Patterns 7, 596–605.

Hamburger, V., and Hamilton, H.L. (1951). A series of normal stages in the development of the chick embryo. J. Morphol. 88, 49–92.

Hare, E.E., Peterson, B.K., Iyer, V.N., Meier, R., and Eisen, M.B. (2008). Sepsid even-skipped enhancers are functionally conserved in Drosophila despite lack of sequence conservation. PLoS Genet. 4, e1000106.

Hu, H., Uesaka, M., Guo, S., Shimai, K., Lu, T.-M., Li, F., Fujimoto, S., Ishikawa, M., Liu, S., Sasagawa, Y., et al. (2017). Constrained vertebrate evolution by pleiotropic genes. Nat. Ecol. Evol. 1, 1722–1730.

Irie, N., and Kuratani, S. (2014). The developmental hourglass model: a predictor of the basic body plan? Development 141, 4649–4655.

Jacob, F. (1977). Evolution and tinkering. Science 196, 1161–1166.

Jung, H., and Dasen, J.S. (2015). Evolution of Patterning Systems and Circuit Elements for Locomotion. Dev. Cell 32, 408–422.

Karlstrom, R.O., Trowe, T., Klostermann, S., Baier, H., Brand, M., Crawford, A.D., Grunewald, B., Haffter, P., Hoffmann, H., Meyer, S.U., et al. (1996). Zebrafish mutations affecting retinotectal axon pathfinding. Development 123, 427–438.

Karlstrom, R.O., Tyurina, O. V., Kawakami, A., Nishioka, N., Talbot, W.S., Sasaki, H., and Schier, A.F. (2003). Genetic analysis of zebrafish gli1 and gli2 reveals divergent requirements for gli genes in vertebrate development. Development 130, 1549–1564.

Ke, Z., Kondrichin, I., Gong, Z., and Korzh, V. (2008). Combined activity of the two Gli2 genes of zebrafish play a major role in Hedgehog signaling during zebrafish neurodevelopment. Mol. Cell. Neurosci. 37, 388–401.

Kiehn, O. (2016). Decoding the organization of spinal circuits that control locomotion. Nat. Rev. Neurosci. 17, 224–238.

Krauss, S., Concordet, J.P., and Ingham, P.W. (1993). A functionally conserved homolog of the Drosophila segment polarity gene hh is expressed in tissues with polarizing activity in zebrafish embryos. Cell 75, 1431–1444.

Kutejova, E., Sasai, N., Shah, A., Gouti, M., and Briscoe, J. (2016). Neural Progenitors Adopt Specific Identities by Directly Repressing All Alternative Progenitor Transcriptional Programs. Dev. Cell 36, 639–653.

Lai, H.C., Seal, R.P., and Johnson, J.E. (2016). Making sense out of spinal cord somatosensory development. Development 143, 3434–3448.

Lee, A.P., Kerk, S.Y., Tan, Y.Y., Brenner, S., and Venkatesh, B. (2011). Ancient Vertebrate Conserved Noncoding Elements Have Been Evolving Rapidly in Teleost Fishes. Mol. Biol. Evol. 28, 1205–1215.

Lewis, K.E., Bates, J., and Eisen, J.S. (2005). Regulation of iro3 expression in the zebrafish spinal cord. Dev. Dyn. 232, 140–148.

Li, S., Misra, K., Matise, M.P., and Xiang, M. (2005). Foxn4 acts synergistically with Mash1 to specify subtype identity of V2 interneurons in the spinal cord. Proc. Natl. Acad. Sci. U. S. A. 102, 10688–10693.

Lu, D.C., Niu, T., and Alaynick, W.A. (2015). Molecular and cellular development of spinal cord locomotor circuitry. Front. Mol. Neurosci. 8, 25.

Ludwig, M.Z., Patel, N.H., and Kreitman, M. (1998). Functional analysis of eve stripe 2 enhancer evolution in Drosophila, rules governing conservation and change. Development 125, 949–958.

Ludwig, M.Z., Bergman, C., Patel, N.H., and Kreitman, M. (2000). Evidence for stabilizing selection in a eukaryotic enhancer element. Nature 403, 564–567.

Luo, J., Ju, M.J., and Redies, C. (2006). Regionalized cadherin-7 expression by radial glia is regulated by Shh and Pax7 during chicken spinal cord development. Neuroscience 142, 1133–1143.

Marillat, V., Sabatier, C., Failli, V., Matsunaga, E., Sotelo, C., Tessier-Lavigne, M., and Chédotal, A. (2004). The slit receptor Rig-1/Robo3 controls midline crossing by hindbrain precerebellar neurons and axons. Neuron 43, 69–79.

Martín-Durán, J.M., Pang, K., Børve, A., Lê, H.S., Furu, A., Cannon, J.T., Jondelius, U., and Hejnol, A. (2018). Convergent evolution of bilaterian nerve cords. Nature 553, 45–50.

Matise, M.P., Epstein, D.J., Park, H.L., Platt, K.A., and Joyner, A.L. (1998). Gli2 is required for induction of floor plate and adjacent cells, but not most ventral neurons in the mouse central nervous system. Development 125, 2759–2770.

Mayran, A., Khetchoumian, K., Hariri, F., Pastinen, T., Gauthier, Y., Balsalobre, A., and Drouin, J. (2018). Pioneer factor Pax7 deploys a stable enhancer repertoire for specification of cell fate. Nat. Genet. 50, 259–269.

McGregor, A.P., Shaw, P.J., Hancock, J.M., Bopp, D., Hediger, M., Wratten, N.S., and Dover, G.A. (2001). Rapid restructuring of bicoid-dependent hunchback promoters within and between Dipteran species: implications for molecular coevolution. Evol. Dev. 3, 397– 407.

Metzis, V., Steinhauser, S., Pakanavicius, E., Gouti, M., Stamataki, D., Ivanovitch, K., Watson, T., Rayon, T., Mousavy Gharavy, S.N., Lovell-Badge, R., et al. (2018). Nervous System Regionalization Entails Axial Allocation before Neural Differentiation. Cell 175, 1105–1118.e17.

Miyamoto, N., and Wada, H. (2013). Hemichordate neurulation and the origin of the neural tube. Nat. Commun. 4, 2713.

Nishi, Y., Zhang, X., Jeong, J., Peterson, K.A., Vedenko, A., Bulyk, M.L., Hide, W.A., and McMahon, A.P. (2015). A direct fate exclusion mechanism by Sonic hedgehog-regulated transcriptional repressors. Development 142, 3286–3293.

Niwa, H., Yamamura, K., and Miyazaki, J. (1991). Efficient selection for high-expression transfectants with a novel eukaryotic vector. Gene 108, 193–199.

Oosterveen, T., Kurdija, S., Alekseenko, Z., Uhde, C.W., Bergsland, M., Sandberg, M., Andersson, E., Dias, J.M., Muhr, J., and Ericson, J. (2012). Mechanistic Differences in the Transcriptional Interpretation of Local and Long-Range Shh Morphogen Signaling. Dev. Cell 23, 1006–1019.

Panayi, H., Panayiotou, E., Orford, M., Genethliou, N., Mean, R., Lapathitis, G., Li, S., Xiang, M., Kessaris, N., Richardson, W.D., et al. (2010). Sox1 is required for the specification of a novel p2-derived interneuron subtype in the mouse ventral spinal cord. J. Neurosci. 30, 12274–12280.

Paris, M., Kaplan, T., Li, X.Y., Villalta, J.E., Lott, S.E., and Eisen, M.B. (2013). Extensive Divergence of Transcription Factor Binding in Drosophila Embryos with Highly Conserved Gene Expression. PLoS Genet. 9, e1003748.

Park, H.C., Mehta, A., Richardson, J.S., and Appel, B. (2002). Olig2 Is Required for Zebrafish Primary Motor Neuron and Oligodendrocyte Development. Dev. Biol. 248, 356– 368.

Park, H.L., Bai, C., Platt, K.A., Matise, M.P., Beeghly, A., Hui, C.C., Nakashima, M., and Joyner, A.L. (2000). Mouse Gli1 mutants are viable but have defects in SHH signaling in combination with a Gli2 mutation. Development 127, 1593–1605.

Peng, C.Y., Yajima, H., Burns, C.E., Zon, L.I., Sisodia, S.S., Pfaff, S.L., and Sharma, K. (2007). Notch and MAML Signaling Drives Scl-Dependent Interneuron Diversity in the Spinal Cord. Neuron 53, 813–827.

Persson, M., Stamataki, D., te Welscher, P., Andersson, E., Böse, J., Rüther, U., Ericson, J., and Briscoe, J. (2002). Dorsal-ventral patterning of the spinal cord requires Gli3 transcriptional repressor activity. Genes Dev. 16, 2865–2878.

Peterson, K.A., Nishi, Y., Ma, W., Vedenko, A., Shokri, L., Zhang, X., McFarlane, M., Baizabal, J.-M., Junker, J.P., van Oudenaarden, A., et al. (2012). Neural-specific Sox2 input and differential Gli-binding affinity provide context and positional information in Shh-directed neural patterning. Genes Dev. 26, 2802–2816.

Prince, V.E., Moens, C.B., Kimmel, C.B., and Ho, R.K. (1998a). Zebrafish hox genes : expression in the hindbrain region of wild-type and mutants of the segmentation gene, valentino. Development 125, 393–406.

Prince, V.E., Joly, L., Ekker, M., and Ho, R.K. (1998b). Zebrafish hox genes: genomic organization and modified colinear expression patterns in the trunk. Development 125, 407– 420.

Rabe, N., Gezelius, H., Vallstedt, A., Memic, F., and Kullander, K. (2009). Netrin-1-dependent spinal interneuron subtypes are required for the formation of left-right alternating locomotor circuitry. J. Neurosci. 29, 15642–15649.

Ren, Q., Zhong, Y., Huang, X., Leung, B., Xing, C., Wang, H., Hu, G., Wang, Y., Shimeld, S.M., and Li, G. (2020). Step-wise evolution of neural patterning by Hedgehog signalling in chordates. Nat. Ecol. Evol. 4, 1247–1255.

Riddle, R.D., Johnson, R.L., Laufer, E., and Tabin, C. (1993). Sonic hedgehog mediates the polarizing activity of the ZPA. Cell 75, 1401–1416.

Roelink, H., Augsburger, A., Heemskerk, J., Korzh, V., Norlin, S., Ruiz i Altaba, A., Tanabe, Y., Placzek, M., Edlund, T., Jessell, T.M., et al. (1994). Floor plate and motor neuron induction by vhh-1, a vertebrate homolog of hedgehog expressed by the notochord. Cell 76, 761–775.

Rutherford, S.L., and Lindquist, S. (1998). Hsp90 as a capacitor for morphological evolution. Nature 396, 336–342.

Sabatier, C., Plump, A.S., Le Ma, Brose, K., Tamada, A., Murakami, F., Lee, E.Y.-H.P., and Tessier-Lavigne, M. (2004). The divergent Robo family protein rig-1/Robo3 is a negative regulator of slit responsiveness required for midline crossing by commissural axons. Cell 117, 157–169.

Sagner, A., and Briscoe, J. (2019). Establishing neuronal diversity in the spinal cord: a time and a place. Development 146, dev182154.

Schäfer, M., Kinzel, D., Neuner, C., Schartl, M., Volff, J.N., and Winkler, C. (2005). Hedgehog and retinoid signalling confines nkx2.2b expression to the lateral floor plate of the zebrafish trunk. Mech. Dev. 122, 43–56.

Schmitz, B., Papan, C., and Campos-Ortega, J.A. (1993). Neurulation in the anterior trunk region of the zebrafish Brachydanio rerio. Roux’s Arch. Dev. Biol. 202, 250–259.

Schoenwolf, G.C., and Smith, J.L. (1990). Mechanisms of neurulation: Traditional viewpoint and recent advances. Development 109, 243–270.

Siegal, M.L., and Bergman, A. (2002). Waddington’s canalization revisited: developmental stability and evolution. Proc. Natl. Acad. Sci. U. S. A. 99, 10528–10532.

Stauber, M., Jäckle, H., and Schmidt-Ott, U. (1999). The anterior determinant bicoid of Drosophila is a derived Hox class 3 gene. Proc. Natl. Acad. Sci. U. S. A. 96, 3786–3789.

Stergachis, A.B., Neph, S., Sandstrom, R., Haugen, E., Reynolds, A.P., Zhang, M., Byron, R., Canfield, T., Stelhing-Sun, S., Lee, K., et al. (2014). Conservation of trans-acting circuitry during mammalian regulatory evolution. Nature 515, 365–370.

Stolfi, A., Lowe, E.K., Racioppi, C., Ristoratore, F., Brown, C.T., Swalla, B.J., and Christiaen, L. (2014). Divergent mechanisms regulate conserved cardiopharyngeal development and gene expression in distantly related ascidians. Elife 3, e03728.

Sun, J., Rockowitz, S., Xie, Q., Ashery-Padan, R., Zheng, D., and Cvekl, A. (2015). Identification of in vivo DNA-binding mechanisms of Pax6 and reconstruction of Pax6-dependent gene regulatory networks during forebrain and lens development. Nucleic Acids Res. 43, 6827–6846.

Swanson, C.I., Schwimmer, D.B., and Barolo, S. (2011). Rapid evolutionary rewiring of a structurally constrained eye enhancer. Curr. Biol. 21, 1186–1196.

True, J.R., and Haag, E.S. (2001). Developmental system drift and flexibility in evolutionary trajectories. Evol. Dev. 3, 109–119.

Tulloch, A.J., Teo, S., Carvajal, B. V., Tessier-Lavigne, M., and Jaworski, A. (2019). Diverse spinal commissural neuron populations revealed by fate mapping and molecular profiling using a novel Robo3Cre mouse. J. Comp. Neurol. 527, 2948–2972.

Tyurina, O. V., Guner, B., Popova, E., Feng, J., Schier, A.F., Kohtz, J.D., and Karlstrom, R.O. (2005). Zebrafish Gli3 functions as both an activator and a repressor in Hedgehog signaling. Dev. Biol. 277, 537–556.

Vanderlaan, G., Tyurina, O. V., Karlstrom, R.O., and Chandrasekhar, A. (2005). Gli function is essential for motor neuron induction in zebrafish. Dev. Biol. 282, 550–570.

Vierstra, J., Rynes, E., Sandstrom, R., Zhang, M., Canfield, T., Hansen, R.S., Stehling-Sun, S., Sabo, P.J., Byron, R., Humbert, R., et al. (2014). Mouse regulatory DNA landscapes reveal global principles of cis-regulatory evolution. Science 346, 1007–1012.

Waddington, C.H. (1942). Canalization of development and the inheritance of acquired characters. Nature 150, 563–565.

Waddington, C.H. (1957). The strategy of the genes (George Allen & Unwin Ltd).

Wang, H., Lei, Q., Oosterveen, T., Ericson, J., and Matise, M.P. (2011). Tcf/lef repressors differentially regulate shh-gli target gene activation thresholds to generate progenitor patterning in the developing CNS. Development 138, 3711–3721.

Wang, X., Zhao, Z., Muller, J., Iyu, A., Khng, A.J., Guccione, E., Ruan, Y., and Ingham, P.W. (2013). Targeted inactivation and identification of targets of the Gli2a transcription factor in the zebrafish. Biol. Open 2, 1203–1213.

Wilson, S.I., Shafer, B., Lee, K.J., and Dodd, J. (2008). A Molecular Program for Contralateral Trajectory: Rig-1 Control by LIM Homeodomain Transcription Factors. Neuron 59, 413–424.

Xiong, F., Tentner, A.R., Huang, P., Gelas, A., Mosaliganti, K.R., Souhait, L., Rannou, N., Swinburne, I.A., Obholzer, N.D., Cowgill, P.D., et al. (2013). Specified neural progenitors sort to form sharp domains after noisy Shh signaling. Cell 153, 550–561.

Zagorski, M., Tabata, Y., Brandenberg, N., Lutolf, M.P., Tkačik, G., Bollenbach, T., Briscoe, J., and Kicheva, A. (2017). Decoding of position in the developing neural tube from antiparallel morphogen gradients. Science 356, 1379–1383.

Zhang, Y., Narayan, S., Geiman, E., Lanuza, G.M., Velasquez, T., Shanks, B., Akay, T., Dyck, J., Pearson, K., Gosgnach, S., et al. (2008). V3 Spinal Neurons Establish a Robust and Balanced Locomotor Rhythm during Walking. Neuron 60, 84–96.

