## Supplemental Information for "The developmental hourglass model is applicable to the spinal cord"

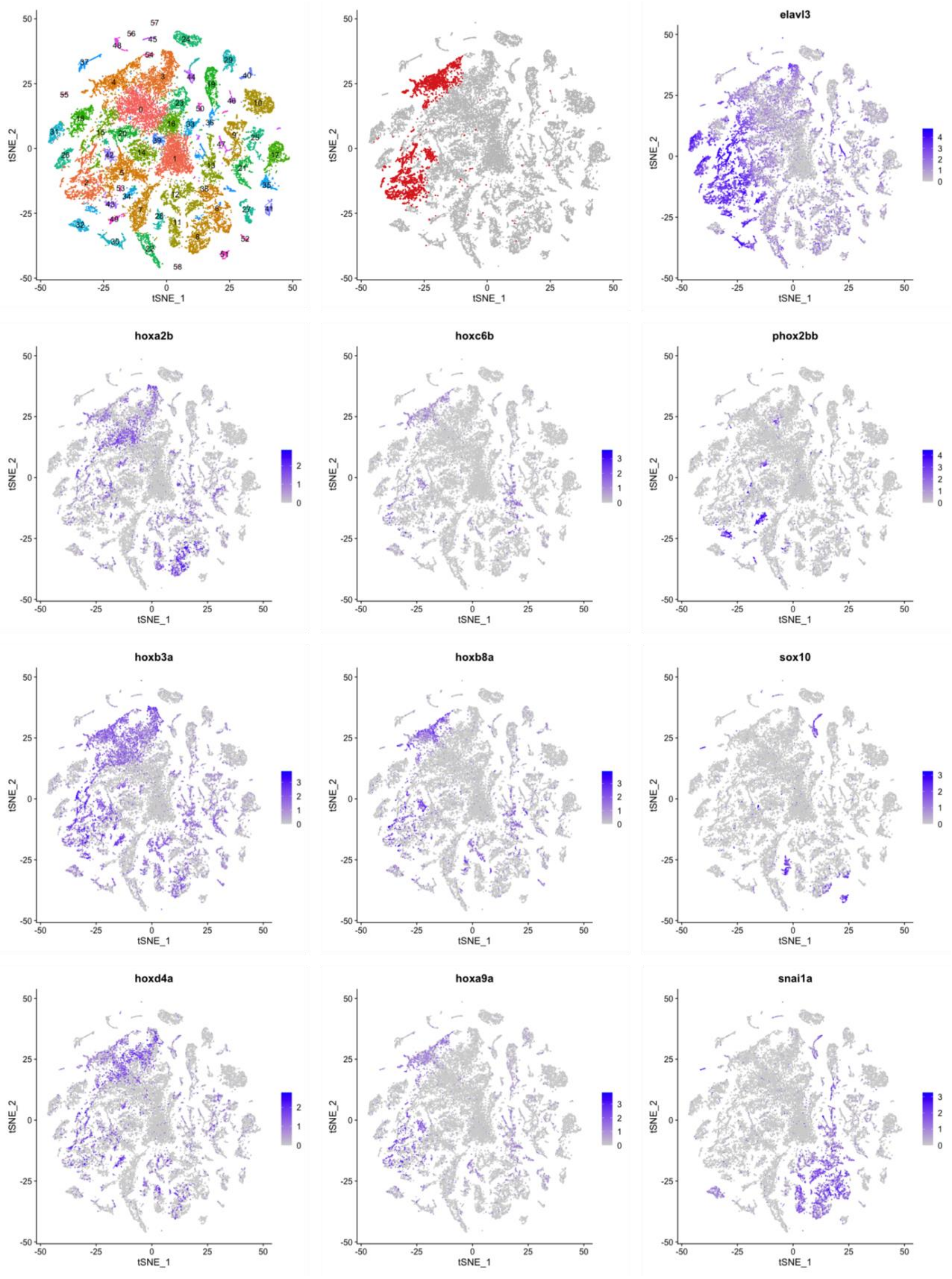

**Figure S1. Subsetting scRNA-seq data derived from zebrafish whole embryo**

scRNA-seq data derived from the zebrafish whole embryos at 1 and 2 dpf were analyzed. Clustering results and gene expressions are visualized on the tSNE plot. Top left panel shows distinct clusters labeled with serial numbers 0–58. To define the spinal cord cells (colored red in the top middle panel), several marker genes and *hox* genes are examined. The expression of *elavl3* roughly indicates the neuronal population. Anterior limit of the expression of 3' *hox* genes, such as *hoxa2b*, *hoxb3a*, and *hoxd4a*, are at the hindbrain (Prince et al., 1998a). Whereas *hoxc6b*, *hoxb8a*, and *hoxa9a* is expressed in the spinal cord, but not in the hindbrain (Prince et al., 1998b), indicating that clusters numbered 0, 3, and 15 are likely to be the hindbrain. We also identified the branchiomotor and cranial parasympathetic preganglionic neurons (*phox2bb*), neural crest cells (*sox10*), and mesodermal cells (*snai1a*) to exclude for the downstream analysis. This allow us to define the cells of the spinal cord, which are cluster numbers 2, 4, and 42.

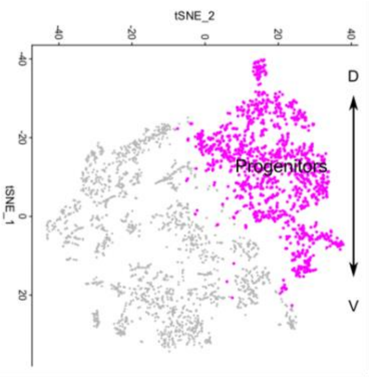
tSNE_2
-40
-20
0
20
40
tSNE_1
D
V
Progenitors

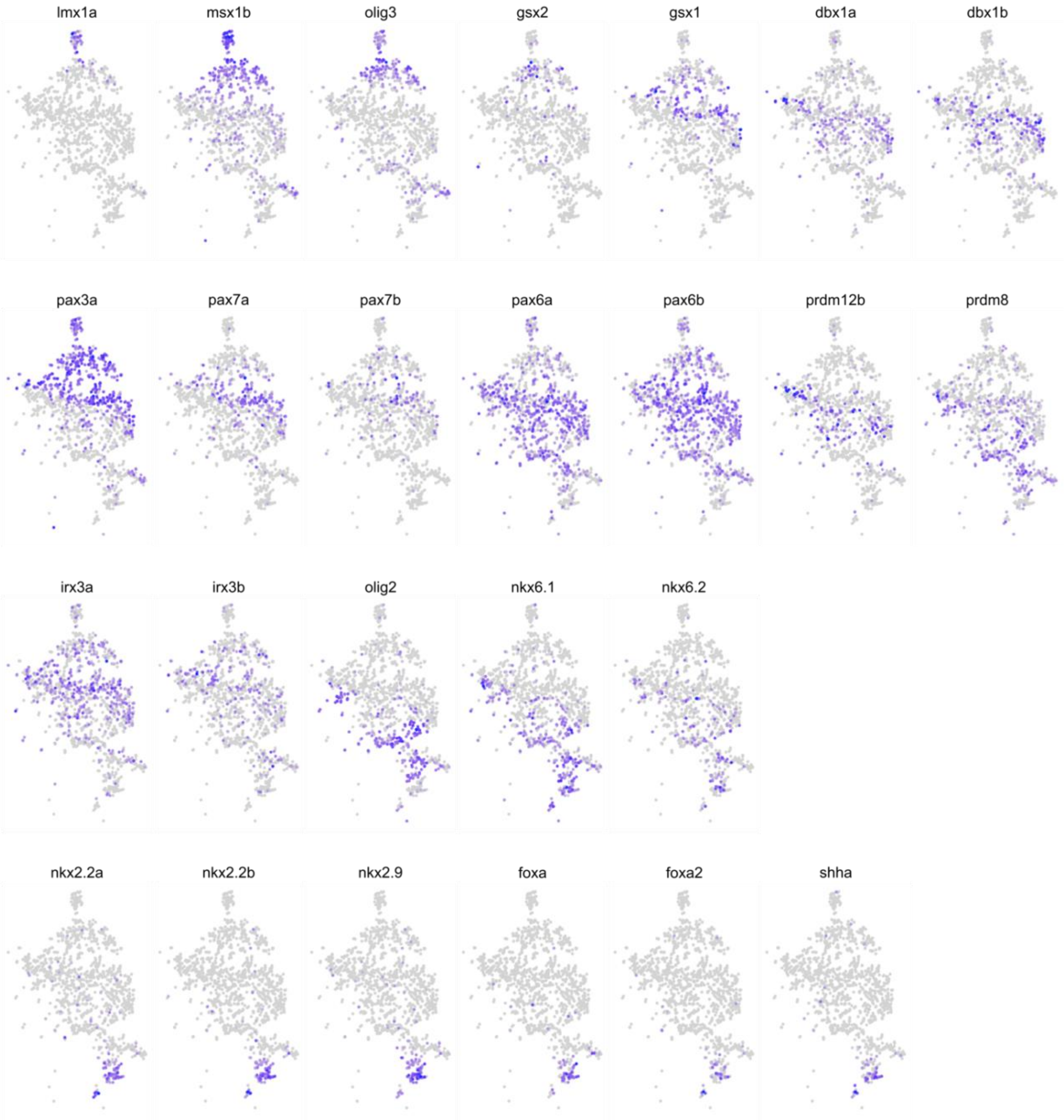
lmx1a
msx1b
olig3
gsx2
gsx1
dbx1a
dbx1b
pax3a
pax7a
pax7b
pax6a
pax6b
prdm12b
prdm8
irx3a
irx3b
olig2
nkx6.1
nkx6.2
nkx2.2a
nkx2.2b
nkx2.9
foxa
foxa2
shha

**Figure S2. Gene expression in the progenitor domains of the zebrafish spinal cord**

Top panel is identical to the tSNE plot in Figure 2, but is rotated 90 degrees clockwise. All other panels show the expression of domain specific TFs. Only progenitor cells are shown. Cell arrangement in this plot is parallel to that *in vivo* along dorsal-ventral axis. D and V indicate dorsal and ventral, respectively. These data support that progenitor domain organization is highly conserved in vertebrates.

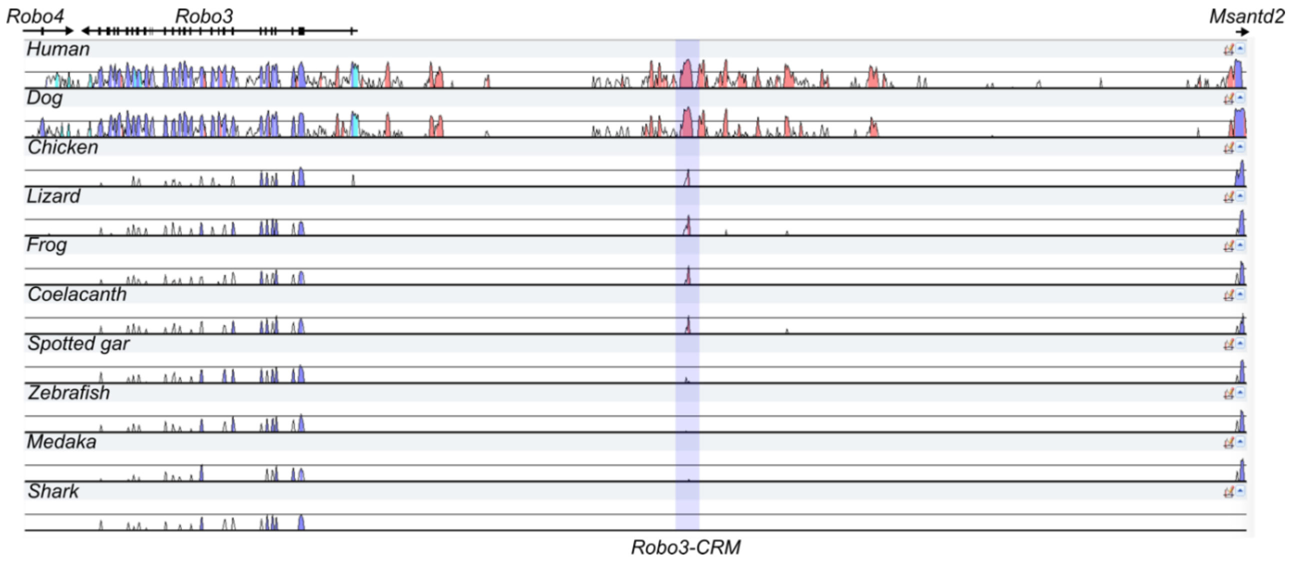


**Figure S3. Alignment of *Robo3* locus visualized by VISTA**

VISTA plot of *Robo3* locus. Genomic region spanning from *Robo3* to the edge of *Msantd2* were aligned. Species compared were indicated in left side. *Robo3-CRM* is highlighted. Guide tree for multiple alignment: (((((((Mouse Human) Dog) (Chicken Lizard)) Frog) Coelacanth) (Spotted_gar (Zebrafish Medaka))) Shark).

Table S2

Sequences of primers used for cDNA amplification for RNA probe preparation

| | |
|---|---|
| ROBO3 Forward | TCCAACTCCTCCGAGCTGCTGCTCGGC |
| ROBO3 Reverse | ATGGAGATGGGCGCACTGCGAGCACCC |
| GSX1 Forward | AGCTGAATTCCTTCCTGGTGGACTCGCTG |
| GSX1 Reverse | GTACAAGCTTCAAAATCATCCGGCTGCACG |
| DBX2 Forward | AGCTGAATTCCAGCCTGGGCAAAAGTTTCC |
| DBX2 Reverse | GTACAAGCTTGCTCTTGGAGGTGGTTGTGT |
| IRX3 Forward | AGCTGAATTCAGTACATCAGGCCGCTGTACCC |
| IRX3 Reverse | GTACAAGCTTTCCCTCTTGTCCTCTTCCCCCT |

Table S3

CRM positions examined in this study

| CRM | Position in mouse reference genome (mm10) |
|---|---|
| Robo3-CRM | chr9:37,453,722-37,454,773 |
| Pax6-CRM | chr2:105,611,719-105,612,835 |
| Gsx1-CRM | chr5:147,155,012-147,156,180 |
| Dbx2-CRM | chr15:95,701,894-95,703,059 |
| Irx3-CRM1 | chr8:91,866,617-91,867,812 |
| Irx3-CRM2 | chr8:91,875,790-91,876,952 |
| Olig2-CRM2 | chr16:91,172,997-91,174,126 |

Table S4
Antibodies used in this study

| Antibody | Source | ID |
|---|---|---|
| Mouse anti-Neurofilament-associated antigen | Developmental studies hybridoma bank (DSHB) | 3A10 |
| Mouse anti-Nkx2-2 | DSHB | 74.5A5 |
| Mouse anti-Lhx1 | DSHB | 4F2 |
| Mouse anti-Isl1/2 | DSHB | 39.4D5 |
| Mouse anti-Evx1/2 | DSHB | 99.1-3A2 |
| Mouse anti-Pax6 | DSHB | PAX6 |
| Rabbit anti-Olig2 | Millipore | AB9610 |
| Mouse anti-GFP | DSHB | DSHB-GFP-4C9 |
| Rabbit anti-GFP | MBL | 598 |
| Rabbit anti-GFP, Alexa Fluor 488 conjugated | Thermo Fisher | A21311 |
| Alexa Fluor 488 anti-mouse IgG1 | Thermo Fisher | A21121 |
| Alexa Fluor 488 anti-mouse IgG2b | Thermo Fisher | A21141 |
| Alexa Fluor 488 anti-rabbit IgG | Thermo Fisher | A11008 |
| Alexa Fluor 594 anti-mouse IgG1 | Thermo Fisher | A21125 |
| Alexa Fluor 647 anti-mouse IgG1 | Thermo Fisher | A21240 |
| Alexa Fluor 647 anti-mouse IgG2b | Thermo Fisher | A21242 |
| Alexa Fluor Plus 647 anti-rabbit IgG | Thermo Fisher | A32733 |
| Goat anti-rabbit IgG, biotinylated | Vector laboratories | BA-1000 |
| Horse anti-mouse IgG, biotinylated | Vector laboratories | BA-2000 |
| Anti-DIG-AP | Sigma-Aldrich | 11093274910 |
